# To close or to collapse: the role of charges on membrane stability upon pore formation

**DOI:** 10.1101/2020.08.31.274860

**Authors:** Rafael B. Lira, Fernanda S.C. Leomil, Renan J. Melo, Karin A. Riske, Rumiana Dimova

**Affiliations:** Departamento de Biofísica, Universidade Federal de São Paulo, São Paulo, Brazil; Department of Theory and Biosystems, Max Planck Institute of Colloids and Interfaces, Potsdam, Germany; Instituto de Física, Universidade de São Paulo, São Paulo, Brazil; Moleculaire Biofysica, Zernike Instituut, Rijksuniversiteit, Groningen, the Netherlands

**Keywords:** giant unilamellar vesicles, charged lipids, edge tension, electroporation, detergent, membrane solubilization, calcium

## Abstract

Resealing of membrane pores is crucial for cell survival. We study membrane surface charge and medium composition as defining regulators of membrane stability. Pores are generated by electric field or detergents. Giant vesicles composed of zwitterionic and negatively charged lipids mixed at varying ratios are subjected to a single strong electric pulse. Charged vesicles are prone to catastrophic collapse transforming them into tubular structures. The spectrum of destabilization responses includes the generation of long-living submicroscopic pores and partial vesicle bursting. The origin of these phenomena is related to the membrane edge tension, which governs pore closure. This edge tension significantly decreases as a function of the fraction of charged lipids. Destabilization of charged vesicles upon pore formation is universal – it is also observed with other poration stimuli. Disruption propensity is enhanced for membranes made of lipids with higher degree of unsaturation. It can be reversed by screening membrane charge in the presence of calcium ions. We interpret the observed findings in light of theories of stability and curvature generation and discuss mechanisms acting in cells to prevent total membrane collapse upon poration. Enhanced membrane stability is crucial for the success of electroporation-based technologies for cancer treatment and gene transfer.

## 1. Introduction

The plasma membrane constitutes the boundaries of cells and acts as an active barrier that separates different compartments and regulates cellular communication and transport. Consisting of a quasi-two-dimensional fluid structure of a few nanometers in thickness, biomembranes can be easily bent but impose a large resistance to stretching. Remarkably, these material properties arise from the physical-chemical characteristics of pure lipid bilayers ^[1]^, and are essential for attributing both mechanical stability and flexibility to cells. When subjected to strong loads, lipid bilayers rupture through the opening of a pore, which tends to eventually reseal and restore membrane integrity as demonstrated on model membranes ^[2]^. Cells are exposed to a range of mechanical stimuli, and even physiological activities may cause membrane disruption leading to pore formation ^[3]^, whereby the healing capacity in nucleated cells depends on Ca^2+^-mediated recruitment of intracellular vesicles ^[4]^. Thus, what makes a cell survive upon membrane damage is its capacity to reseal the formed pores. A large number of therapeutic substances have their target in the cell interior but the barrier function of the membrane prevents their entry. Therefore, biotechnological applications for intracellular delivery make use of approaches relying on pore formation in the plasma membrane ^[5]^. One of them employs membrane electroporation allowing for efficient delivery of materials, based on the application of one or more electric pulses. Electroporation-based technologies extend to a growing number of applications including cancer treatment by electrochemotherapy ^[6]^, genetic transformation of microorganisms as in gene therapy and DNA vaccination ^[7]^, treatment of cardiac tissue and tumor ablation ^[8]^, and microorganism inactivation as in waste-water treatment and food pasteurization ^[9]^. All of these technologies rely on the formation of pores in the cell membranes. It remains a mystery however, why these pores have a wide range of lifetimes that can span from milliseconds to minutes ^[10]^ (or may not even close at all). Pore stability depends on experimental factors such as the poration parameters, as well as on cellular characteristics such as cell type and anchorage to the cytoskeleton ^[11]^. Alternatively, when induced by chemical agents such as detergents or pore-forming toxins, the stability of pores depends on the way these molecules interact with the membrane. Because of the relation to cell physiology and biotechnology, understanding membrane pore formation and, more importantly, plasma membrane stability is of fundamental and crucial significance.

Model membranes have been widely used to characterize pore dynamics averting the influence of underlying active cell processes. Among all systems, giant unilamellar vesicles (GUVs) are perhaps the best suited model as they represent a free-standing bilayer virtually devoid of artifacts ^[12]^. These vesicles are large enough to be directly visualized and manipulated under an optical microscope displaying the membrane response to external perturbations ^[13]^. Pore formation in GUVs can be indirectly detected through membrane permeability assays or directly visualized in the case of optically resolved pore sizes. Pores in GUVs can be induced by a variety of chemical and mechanical stimuli, including antimicrobial peptides ^[14]^, pore-forming proteins ^[15]^, detergents ^[16]^, photochemical reactions ^[2a]^, membrane stretching ^[17]^ and electroporation ^[2b, 18]^. Many studies have focused on the mechanisms of pore creation, while often neglecting pore stability and closure pathway. However, it is the ability of the formed pores to close that dictates the fate of the cell in its struggle to survive. The lifetime of artificially-triggered pores also defines the efficiency of the above-mentioned biotechnology approaches requiring transport through the membrane. Electric pulses as means of porating vesicles are convenient as their impact is easy to modulate and focuses on the vesicle poles or cell surface facing the electrodes. Electroporation of single-component zwitterionic lipid GUVs is relatively well studied as reviewed in Refs. ^[19]^. Because lipid membranes act as insulating shells, the application of electric field results in vesicle deformation, whereby the type of deformation depends on the ratio of the solution conductivity across the membrane ^[20]^. Above a certain potential threshold, a single direct current (DC) pulse (e.g. 100 µs long) can trigger membrane rupture generating membrane pore(s). In phosphatidylcholine membranes, pulse-triggered optically detected macropores (0.5 – 5 µm in diameter) have lifetimes on the order of 50 ms ^[2b, 21]^. The dynamics of the pores in a lipid bilayer is well described by a balance between membrane tension (σ), which tends to open and stabilize the pores, and edge tension (*γ*), which acts to shrink and close the pores ^[22]^. Whereas the former can vary in response to different external stimuli, the latter is an intrinsic material property ^[18, 23]^. When a porating DC pulse is applied to a GUV, pore formation can relieve the surface tension (unless it is externally sustained e.g. by a micropipette ^[18]^) and the formed pores close solely driven by the edge tension.

Importantly, the edge tension is modulated by the membrane composition, and therefore pore dynamics and membrane stability may be modulated accordingly. The bulk of studies targeting membrane/pore dynamics in GUVs have been predominantly performed on single-component vesicles made of zwitterionic lipids, which represent a rather simple cell membrane model. Considering that cancer cells differ from normal ones by exhibiting abnormal negative surface charge ^[24]^, we investigated the impact of charge on the stability of model membranes. Indeed, when electric pulses are applied to charged multicomponent GUVs, the dynamics of electroporation can be dramatically altered as briefly reported in ^[25]^. As we show here, the response can span from long-living submicroscopic pores, to partial vesicle bursting or complete collapse. We explore the vesicle stability as a function of membrane charge. Poration was triggered by electric fields or caused by a detergent. The fraction of destabilized vesicles and the membrane edge tension, which governs pore closure, are assessed as a function of membrane composition, lipid charge and degree of unsaturation, as well as screening in the presence of salts. We interpret the observed findings in light of existing theories and curvature stabilization and discuss mechanisms that may act to prevent full collapse of living cells upon poration.

## 2. Results

We studied the stability of GUVs prepared from mixtures of zwitterionic and anionic lipids as a function of membrane composition (molar fraction of charged lipid and lipid tail unsaturation) and presence of different ion additives in the medium. The lipid compositions consisted of mixtures of palmitoyl-oleoyl phosphatidylcholine (POPC), palmitoyl-oleoyl phosphatidylglycerol (POPG), dioleoyl phosphatidylcholine (DOPC) and dioleoyl phosphatidylglycerol (DOPG). GUVs were prepared in sucrose and dispersed in glucose, see Materials and Methods. The difference in the refractive indexes of these solutions enhances the image contrast of vesicles observed under phase contrast microscopy and allows optical detection of macropores visualized as a disruption of the bright halo around the vesicle (see e.g. Fig. 1).

**FIGURE 1.**
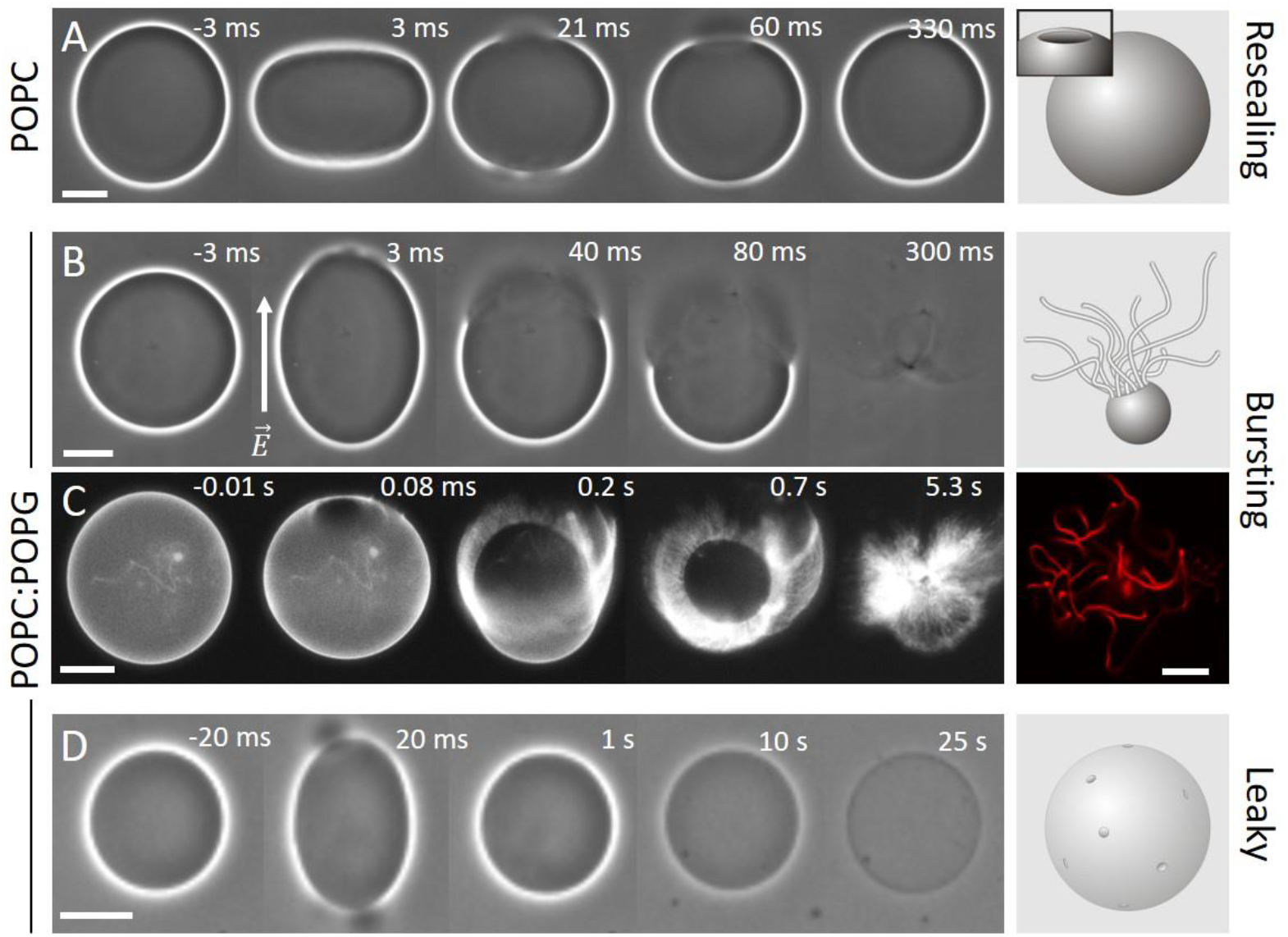
Response of neutral (A, POPC) and charged (B-D, POPC:POPG 1:1) GUVs exposed to a DC pulse (3 kV/cm, 150 µs): (A) macropores reseal fully, (B, C) vesicles burst and collapse into tubular network, and (D) resealing of macropores leaves a leaky vesicle which loses optical contrast (imparted by sugar asymmetry) over time. The time relative to the beginning of the pulse is shown on each snapshot. The field direction is indicated in (B). The images were obtained with phase contrast (A, B, D) or epifluorescence microscopy (C), in which case the membrane contains 0.1 mol% DPPE-Rh. In (A) and (C), the external media contains 0.1 mM NaCl, resulting in oblate deformation of the GUVs upon applying the pulse, whereas in (B) and (D), no salt is present and the vesicles deform into prolate shapes. Scale bars: 10 µm. The sketches on the right illustrate the macropore (A) and the GUV seconds (A, D) or tens of milliseconds (B) after applying the pulse. A confocal image of tubular lipid structures as remnants from the bursting GUV in (C) is also shown on the right.

### 2.1. Electroporation of neutral *versus* charged GUVs: charged membranes are less stable

GUVs were exposed to single DC square-wave pulses of 3 kV/cm magnitude and 150 µs duration. Such pulses are strong enough to induce electroporation in vesicles of radii above ∼7 µm under the conditions explored here, see Text S1 in the Supporting Information (SI). The response of GUVs made of mixtures of POPC and POPG to electric pulses was followed with optical microscopy. If not mentioned otherwise, vesicles prepared from pure POPC and equimolar mixtures of POPC:POPG will be referred to as neutral and negative, respectively.

Upon the application of a single DC pulse, neutral GUVs deform and the membrane ruptures forming one or more visible macropores of few micrometers in diameter. A typical sequence of vesicle deformation, poration and macropore closure of a neutral GUV upon pulse application is shown in Fig. 1A. Initially, the vesicle deforms and visible macropores are formed (at 21-60 ms). Then, all formed pores close, the vesicle relaxes back to its initial spherical shape and the membrane integrity is restored. In contrast, when similar pulses are applied to negatively charged vesicles at identical conditions, they frequently burst through continuous pore expansion leading to complete vesicle collapse (Fig. 1B and Movie S1 in the SI). The fraction of GUVs that undergo bursting depends on membrane composition, as we will discuss below.

Fluorescence microscopy reveals that pore expansion in the charged vesicles occurs on the expense of transforming the locally quasi-flat membrane of the GUV into highly curved structures such as buds and tubes initiating from the pore rim (Fig. 1C, Fig. S1A and Movie S2). Vesicle bursting is a very fast process and completes within a few hundred milliseconds. The internal content of the GUVs is rapidly released (Fig. S1 and Movie S3). In some cases, part of the GUV membrane is still able to close back to a smaller vesicle connected to many tubules present in the region where the macropore closed (Fig. S2); we will refer to this process as partial bursting. The excessive tubulation indicates that charged lipids may be involved in modulating membrane curvature and that this curvature is not entirely relaxed upon membrane poration. We evaluated the fluorescence intensity of the tubes formed after partial or complete bursting of charged vesicles, as an indirect measure of the tube diameter (Fig. S3). The tubes are progressively thinner for increasing charge density. Remarkably, tubes formed upon complete bursting of GUVs containing 50 mol% PG are thinner than tubes that are formed on GUVs that did not collapse (partial bursting), suggesting the role of membrane spontaneous curvature in defining the stability and fate of an electroporated membrane. The mechanism of curvature generation is presumably related to bilayer lipid asymmetry ^[26]^ in our system, also present in cell membranes emphasizing the relevance of the observations.

The experiments above suggest that the pore evolution depends on membrane charge. It is known that some molecules can preferentially accumulate along the pore edges stabilizing them, see e.g. ^[27]^. We thus attempted to assess differences between the lipid composition along the pore rim and regions of intact membrane by using fluorescent probes with different characteristics (i.e. charge, geometry, see Fig. S4A). For all compositions tested and in the presence of the different additives, no dye accumulation or depletion could be detected (Fig. S4B). Considering the short pore lifetime (50-100 ms) and lipid diffusion coefficient of ∼5 µm^2^/s in these membranes ^[28]^, dye accumulation may not be detected simply because the pores close too fast. Even in instances of long-living macropores (∼2 s), no dye accumulation/depletion was detected (Fig. S4C and Movie S4). To slow down pore closure to a couple of seconds to minutes, we electroporated GUVs embedded in an agarose gel ^[21, 28b]^ but no dye accumulation was observed, Fig. S5 and Movie S5. We conclude that in the tested experimental conditions, the composition along the pore rim is similar to the composition in regions of intact membranes regardless of overall membrane charge or probe type.

### 2.2. Submicroscopic pores persist after electro-poration rendering the vesicles leaky

While bursting (partial or complete) is observed exclusively on negatively charged and not on neutral GUVs, only a fraction of the negative vesicles undergoes bursting. In the remaining population, formed pores reseal similarly to those in neutral GUVs. However, some of the negative GUVs that survive macroporation lose their optical contrast seconds after the end of the pulse, see Fig. 1D. This indicates that (at least temporarily) the membrane remains leaky: the original sugar asymmetry vanishes and so does the optical contrast. Figure 2A shows the relative optical contrast in vesicles of different composition as a function of time after applying the electric pulse (see Fig. S6 for details on measurements). For GUVs with long-term permeability, the contrast loss completes within 1-2 minutes in an all-or-none fashion. Increasing the fraction of charged lipids does not seem to alter the rate of permeation but increases the fraction of vesicles with long-term permeability.

**FIGURE 2.**
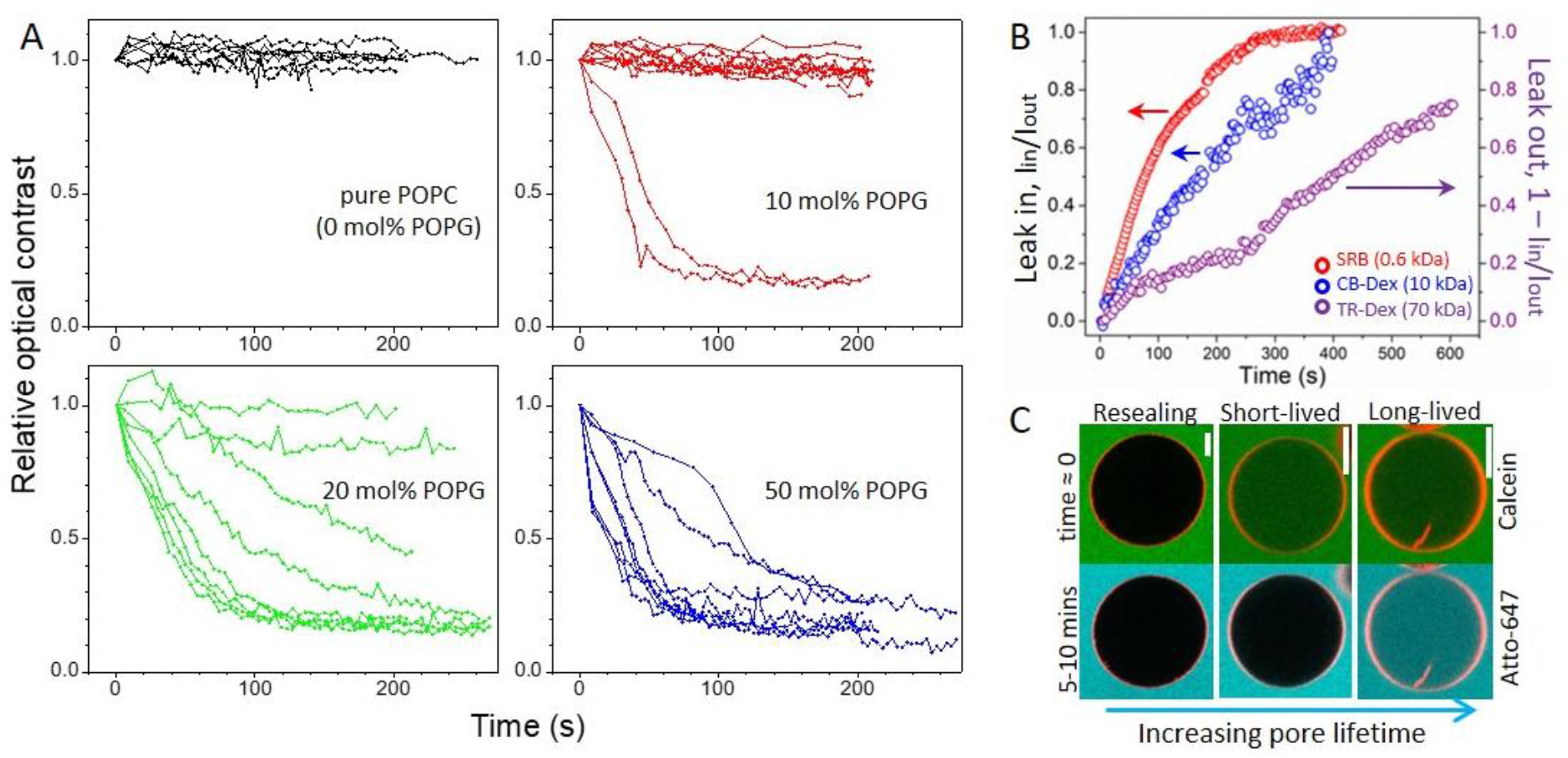
Charged GUVs surviving electroporation remain in a highly permeable state after the pulse (3 kV/cm, 150 µs). (A) Time dependence of the relative contrast loss for GUVs made of POPC with increasing molar fraction of POPG. Each curve represents a measurement on a single GUV (8-10 vesicles per composition were measured). Time 0 indicates when the pulse was applied. To evaluate the contrast loss (see Fig. S6), data were normalized by the value obtained right after macropore closure and relaxation of the vesicle to a spherical shape (typically 50-100 ms after the pulse). (B) Long term permeation of the water-soluble dyes SRB (red) and CB-Dex (blue) into 50 mol% POPG GUV after electroporation shown as the fluorescence intensity ratio I_in_/I_out_ measured inside/outside the same vesicle. Data in purple show the slower leak out (1 – I_in_/I_out_) of TR-Dex from another GUV. (C) Three types of vesicle response (membrane in red; 50 mol% POPG) illustrated with dye entry at different times after electroporation. Calcein (green) is added prior to poration (time ≈ 0, upper row) and shows the immediate dye permeation occurring in the first 1-2 minutes after the pulse as well as long-term permeability. Atto-647 (cyan, lower row) is added 5-10 minutes after electroporation and reports the presence of long-lived pores only. Scale bars: 20 µm.

To roughly assess the minimum size of the long-lived pores (not resolved optically), we performed electroporation in the presence of dyes of increasing molecular weight: sulforhodamine B (SRB, 0.6 kDa), cascade blue-labeled dextran 10 kDa (CB-Dex, 10 kDa) and Texas red-labeled dextran 70 kDa (TR-Dex, 70 kDa) with respective hydrodynamic diameters of approximately 1, 4 and 13 nm ^[29]^. The first two dyes were added in the external medium, whereas TR-Dex was encapsulated inside the GUVs to avoid adhesion to the glass chamber. The entrance of dyes into the GUVs while the macropore is open is negligible (∼1% signal increase for both neutral and negative GUVs, Fig. S7) because of the short pore lifetime. Figure 2B shows the permeation of SRB and CB-Dex into a single negative GUV quantified by the ratio of internal (I_in_) to external fluorescence (I_out_) intensities over time. Both dyes permeate completely until full equilibration across the membrane, whereby the smaller dye (SRB) is faster, as expected. Purple data show the release (leak out) of TR-Dex from a negative GUV, expressed as (1-I_in_/I_out_). Even slower permeation of TR-Dex is observed, as expected for its larger size. More importantly, these results show that the persisting nanopores on negative GUVs are larger than 13 nm.

We then tested whether this highly permeable state persisted minutes after pulse application. Negative GUVs were electroporated in the presence of calcein; such small dyes (< 1 nm) reach concentration equilibrium across the vesicle membrane already within ∼2 min (see Fig. 2A,B). Approximately 5-10 minutes later, a second dye, Atto-647 (< 1 nm) was added to the chamber. Figure 2C shows the three observed outcomes: (i) When the GUVs do not become permeable to calcein, they also exclude Atto-647, and these correspond to the most stable, fully resealed vesicles. (ii) GUVs partially permeable to calcein exclude Atto-647, indicating that pores are short-lived with a delayed closure (less than a couple of minutes). (iii) Vesicles are permeable to both calcein and Atto-647 and thus with long-lived pores or leaky. From three different preparations (43 measured GUVs in total), 37% of the GUVs that survived bursting showed short- and long-lived pores (permeation to calcein), whereas the fraction of vesicles with long-lived pores was 21% (permeation to Atto-647); in other words about half of the GUVs that survived but showed permeation after the pulse, remain permanently permeable. In summary, while the membrane of neutral vesicles reseals completely after poration (Fig. 1A), negatively charged membranes fail to do so and remain perforated for a long time, analogously to pores formed on cells.

### 2.3. Vesicle stability and increased permeability de-pends on fraction of charged lipids, chain unsatura-tion and medium composition

Taken together, the above results demonstrate that charged membranes are less stable when exposed to electric fields and that electroporation results either in: (i) complete vesicle collapse (bursting) and restructuring to membrane nanotubes, (ii) partial bursting leading to a smaller vesicle whereby the membrane around the pore rim transforms into tubes, (iii) macropore closure but maintenance of a permeable state with long-lived sub-microscopic pores (>13 nm in size), or still, (iv) complete membrane resealing.

To quantify how surface charge density affects membrane destabilization, for increasing fraction of POPG lipids in POPC membranes we measured the fraction of vesicles that underwent bursting (X_burst_) as in Fig. 1B,C and the fraction of vesicles in which macropores resealed but GUVs remained in a highly permeable state (X_perm_) as in Fig. 1D. The former represents the fraction of vesicles that burst over the entire population subjected to electroporation: X_burst_ = n_burst_/n_GUVs_, whereas the latter corresponds to the fraction of GUVs that survive bursting (including instances of partial bursting) but lose their contrast: X_perm_ = n_perm_/(n_GUVs_ – n_burst_). Both X_burst_ and X_perm_ increase in a non-trivial manner with increasing the surface charge, i.e., the molar fractions of POPG (see Fig. S8A). We also explored the role of acyl chain unsaturation on membrane destabilization by measuring X_burst_ and X_perm_ for GUVs of dioleoyl (DO) lipids with an unsaturated bond in each of the chains (DOPC:DOPG mixtures) as opposed to saturation in one chain in palmitoyl-oleoyl (PO) lipids (POPC:POPG). As shown in Fig. S8B, contrary to the less unsaturated POPC:POPG vesicles, nearly all DOPC:DOPG vesicles burst at 20 mol% PG or above, and all of the few that survived were permeable, suggesting that higher degree of lipid unsaturation leads to lower membrane stability.

Because GUV bursting and highly permeable states are intimately related to membrane integrity, we combine the two contributions into a fraction of compromised vesicles, X_comp_=(n_burst_+n_perm_)/n_GUVs_. Figure 3A shows X_comp_ as a function of the molar fraction of charged lipid for PO and DO-based mixtures. Even though the absolute values of the fractions varied for different preparations (visualized by the shaded area in Fig. 3A), the overall trend is obvious. Membranes become more unstable upon electroporation as the fraction of charged lipids increases. At each molar fraction of the charged lipid, X_comp_ is higher for DO-based GUVs, confirming our conclusion for lower stability of these membranes. It is important to mention that for the observed behavior, there is no dependence on GUV size (as long as vesicles were visibly porated) or on the proximity to the electrodes, ruling out possible effects of field inhomogeneity. Moreover, as the experiments were performed with fresh solutions, there are no significant changes in media conductivity or aging. In addition, we do not expect membrane destabilization to result from pulse-induced lipid oxidation, as the latter can be observed only after minutes following the pulse and for specific conditions not present in this study ^[30]^. We therefore conclude that increasing charge renders the membranes unstable upon electroporation and this effect is enhanced with the degree of lipid unsaturation.

**FIGURE 3.**
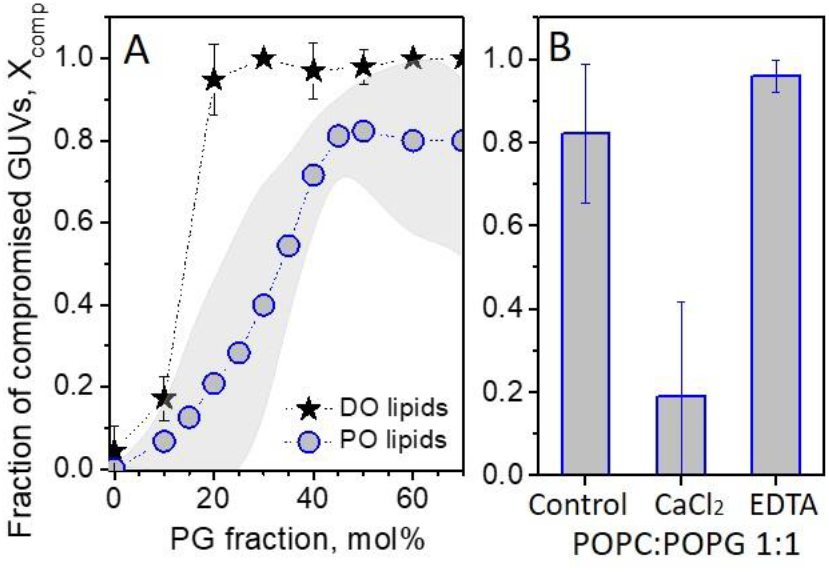
Effect of membrane composition (lipid charge and unsaturation) and environmental conditions on GUVs stability upon electroporation. (A) Fraction of compromised GUVs, X_comp_, made of POPC and DOPC with increasing molar fractions of POPG and DOPG, respectively. For DOPC:DOPG membranes (stars), average values and standard deviations for a number of measurements (> 15 GUVs) are shown for one vesicle preparation per composition; for POPC:POPG membranes (open circles) the gray band illustrates the error from measurements on several preparations per composition. (B) X_comp_ for POPC:POPG (1:1) GUVs electroporated in the presence of 0.5 mM CaCl_2_ or 0.5 mM EDTA. The control measurement corresponds to the standard conditions also used in (A) with 0.1 mM NaCl present in the vesicle exterior.

Next, we studied the effects of certain additives (salts and the calcium chelator EDTA, the respective solution conductivities are given in Table S1) on destabilization of POPC:POPG 1:1 vesicles. Because it is difficult to produce highly charged GUVs in high ionic strength media, the compounds were added exclusively in the external medium. Only low salt concentrations were explored to avoid sample heating and electrolysis upon pulse application at high ionic strength. The addition of 1 mM NaCl in the GUV exterior did not result in differences in the vesicle response (compared to the data in Fig. 3A which was collected for 0.1 mM NaCl). We then explored the effect of CaCl_2_, as calcium ions have higher affinity to charged membranes, see e.g. ^[31]^, and hence different charge neutralization capacity. EDTA was used to remove possible Ca^2+^ contaminants present in the medium ^[32]^, considering that lipid concentration in the samples may be comparable to that of impurities ^[33]^. Figure 3B shows X_comp_ for the conditions tested. In the presence of only 0.5 mM CaCl_2_, X_comp_ is strongly decreased to around 20% whereas, in the presence of 0.5 mM EDTA, X_comp_ increases to nearly 100% (vesicles in the presence of 0.2 mM EDTA have similar behavior, data not shown). In summary, these results show that modest increase in the ionic strength does not alter the membrane behavior while calcium binding stabilizes the membranes. Addition of EDTA, which removes possible divalent ion contaminants from the medium, has the opposite effect.

### 2.4. Membrane edge tension

Pore dynamics and stability are governed by a balance between the edge tension (γ), a material property representing the energy penalty per unit length for arranging the lipids at the pore, and the membrane surface tension (σ), which reflects the stress a membrane is subjected to. Because vesicle bursting is a consequence of continued pore opening, and since the membrane tension is released once the membrane ruptures and the pulse ends, we hypothesized that the increased fraction of compromised vesicles results from decreasing edge tension in charged membranes. We measured γ from the dynamics of macropore closure upon electroporation ^[23b]^, see SI Text S2 and Fig. S9 for details on the method. The experiments were performed with 0.1 or 0.5 mM NaCl at which no measurable effects on X_comp_ were observed (Fig. 3B, control). Figure 4A shows typical pore closure dynamics in GUVs of POPC with increasing POPG fraction. Note the changes in the slope resulting from changes in *γ*. Figure 4B shows a compilation of data measured for GUVs of all tested compositions and Table S2 in the SI lists their mean values. GUVs of pure POPC or containing up to 30 mol% POPG have pore edge tension ∼40 pN (Fig. 4B) comparable to literature data for similar membrane compositions ^[16, 23a, 23b, 34]^. GUVs with 50 mol% POPG show a significant reduction in γ to ∼20 pN. Because the edge tension is determined from pore closure, the measurements are not possible in cases of GUV bursting. The fraction of bursting vesicles is higher for higher fraction of POPG (Fig. 3A), and thus, the mean value of γ is likely to be overestimated at higher POPG fractions. In fact, the lowest γ values shown for 50 mol% POPG arise from cases of partial bursting where very long pore lifetimes are observed (indicated in magenta in Fig. 4B). This difference is exemplified in Fig. S9. Partial bursting corresponds to the lowest possible γ measured and cases of complete bursting are likely to be a result of even lower edge tension values.

**FIGURE 4.**
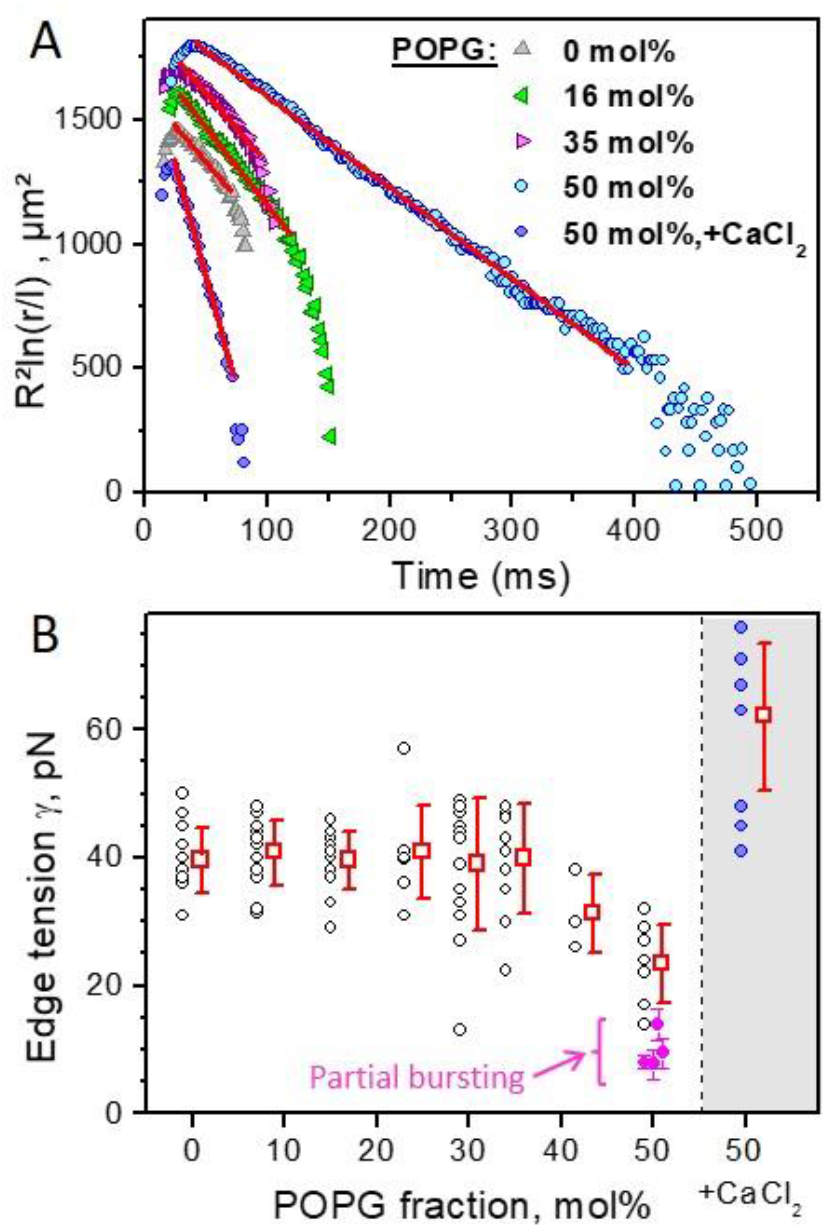
Measurements of edge tension of POPC:POPG GUVs with varied surface charge and under different experimental conditions. (A) Traces of pore closure dynamics in POPC GUVs with increasing fraction of POPG (R is the vesicle radius, r is pore size in µm rescaled by l = 1 µm). The data in the slow pore closure regime are fitted by a linear dependence (red line) the slope of which is used to calculate the edge tension (see text for details). Differences in the absolute values of the data represent mainly differences in GUV size. (B) Edge tension values for increasing fractions of POPG (each point represents a measurement on one vesicle) in the presence of 0.1 or 0.5 mM NaCl (see also SI Table S2). Data in the gray region correspond to measurements on POC:POPG 1:1 GUVs in the presence of 1 mM CaCl_2_. Data collected on vesicles exhibiting partial bursting is also shown (magenta), see Text S2 in the SI.

Next, we tested the effect of CaCl_2_ (which significantly increased the vesicle stability, Fig. 3B) on the edge tension of GUVs of POPC:POPG (1:1), Fig. 4B (shaded area). The presence of 1 mM CaCl_2_ causes a significant increase in the edge tension, to γ = 62 ± 11 pN. Presumably, Ca^2+^ binds and screens the membrane charges, increasing the pore edge tension and stabilizing charged membranes exposed to electroporation. No dependence of the edge tension on the maximal pore size for any composition and regardless of presence of CaCl_2_ was observed (Fig. S10).

In summary, the pore edge tension decreases with increasing the surface charge of the membrane, which is consistent with the increasing fraction of compromised vesicles, X_comp_. The presence of Ca^2+^ increases the pore edge tension and stabilized the vesicles.

### 2.4. Bursting of negative GUVs does not depend of the poration method

The experiments above demonstrate decreased stability of negative membranes upon electroporation. Macroscopic pores in GUVs can be induced upon tension increase imposed also by hypotonic assault ^[35]^, light irradiation ^[2a, 23a]^, adhesion ^[36]^, or chemically, employing e.g. detergents ^[16]^ or proteins such as talin ^[37]^. We questioned whether bursting of charged vesicles depends on the way the macropore is triggered. To rule out the influence of the electric field, we induced pores using the detergent Triton X-100. Consistent with an earlier study ^[38]^, incubation of neutral GUVs with Triton X-100 results in GUV area increase as detergents insert into the membrane, loss of contrast because of membrane permeabilization and eventually full solubilization (Fig. 5A). In the time course of solubilization, macropores are formed and stabilized as a result of detergent-induced decrease in pore edge tension as previously reported ^[16]^. The whole process depends on detergent concentration but typically takes a few minutes and the macropores are stable for dozens of seconds.

**FIGURE 5.**
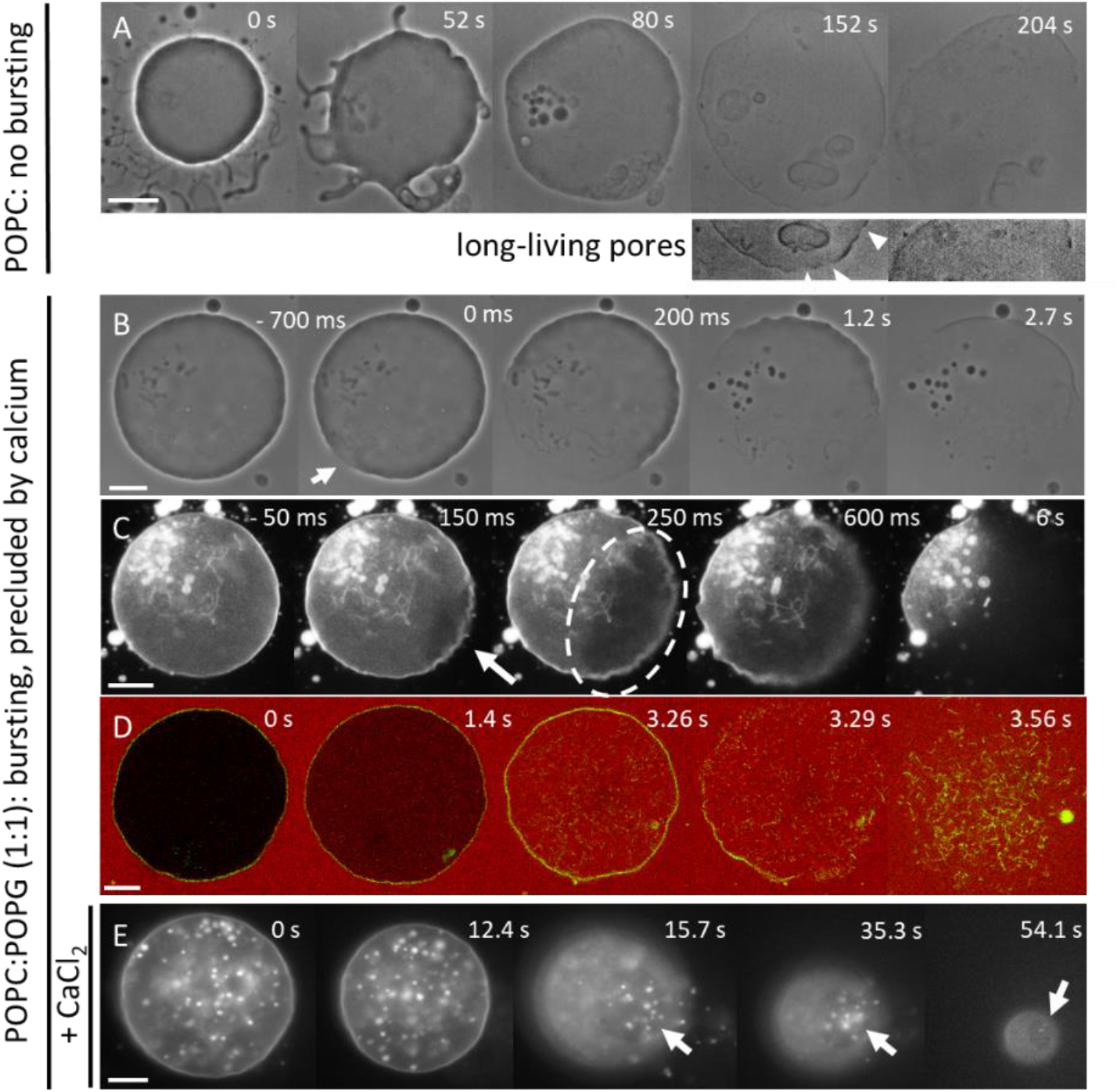
Differential effects of detergent-induced pores in neutral vs. charged GUVs in the presence Triton X-100 at final concentration of 1 mM. (A) On neutral POPC GUVs, Triton X-100 induces large area increase and formation of visible pores that last dozens of seconds before the vesicle is slowly solubilized (see arrowheads in the enhanced cropped images in the second row). (B-D) In sharp contrast, on negative POPC:POPG GUVs the formation and rapid expansion of a single macropore (see arrows and encircled pore region in the middle image in C) proceeds to vesicle bursting strongly resembling response to electroporation. The process is viewed under (B) phase contrast, this sequence is shown in Movie S6; (C) epifluorescence, Movie S7; (D) confocal cross-sections where the outside medium contains 2.5 µM SRB, Movie S8. (E) In the presence of CaCl_2_ (3.5 mM), the macropore (indicated with arrows) stays open during the whole solubilization process, Movie S9. The GUVs in the fluorescence images contain 0.5 mol% DPPE-Rh. All time stamps correspond to the beginning of observation. Scale bars: 20 µm.

The behavior of negatively charged vesicles is strikingly different. The GUVs are initially impermeable to small molecules such as sugars and SRB dispersed in the external medium (see first snapshots in Fig. 5B,D). Upon detergent insertion, the vesicle area increases as usual. However, the stochastic appearance of the first visible macropore results in instantaneous vesicle bursting completing in a few hundreds of milliseconds (Fig. 5B-D, see also Movies S6 and S7) in a way seemingly identical to electroporation. Detergent-induced bursting is also associated with the formation of membrane tubes (Fig. 5D) that are completely solubilized later. Unexpectedly, charged GUVs become permeable to small dyes (< 1 nm) before bursting (see Fig. 5C and Movie S8). This implies that submicron pores are formed before a visible macropore is observed, and only after it becomes large enough does the vesicle burst. In other words, there may be a threshold in pore size for bursting, below which the vesicle is still stable.

We next investigated the detergent-induced poration in the presence of CaCl_2_ to test for possible charge screening effects on GUV bursting as observed during electroporation (Fig. 3B). Initially, the GUV area increases as usual, which in this case is observed as an increase in intravesicular structures (rather than increase in vesicle size). This inward tubulation is likely due to calcium-induced change in membrane spontaneous curvature ^[39]^. The GUV then becomes permeable to small molecules followed by the stochastic opening of a single macropore, through which internal structures can leave the vesicle (Fig. 5E, Movie S9). The macropore remains open for tens of seconds until the whole vesicle is solubilized. This sequence of solubilization resembles that for neutral POPC GUVs, which is not surprising considering the observed stabilization of charged vesicles against bursting ascribed by calcium (Fig. 3). We conclude that both bursting of charged vesicles and calcium-induced stabilization due to charge screening are universal effects and do not depend on the way pores are triggered.

## 3. Discussion

The response of PC:PG GUVs to electric pulses was investigated to shed light on the mechanisms triggering the morphological transformations and membrane destabilization. Considering that contrary to healthy cells, cancer cell typically expose negatively charged lipids, we investigated the charge dependence of the membrane response to resolve whether surface charge is an important cue (e.g. in electrochemotherapeutic approaches). We find pronounced vesicle destabilization with increasing surface charge (Figs. 3 and 5), whereas the edge tension drops down (Fig. 4). Full collapse of macroporated vesicles represents the most extreme destabilization response of charged GUVs. Importantly, bursting is not limited to pores induced by an electric field but seems to be a rather universal response of charged membranes to macropore formation. The frequency of membrane destabilization increases for raising membrane charge density and unsaturation of the acyl chains, and can be reversed in the presence of calcium in the medium (Figs. 3 and 5E).

The material properties of charged and neutral membranes differ. Compared to neutral ones, charged membranes are stiffer ^[40]^, and display reduced lysis tension (and are thus easier to porate) ^[41]^. We show that, in addition to these properties, charged membranes display lower pore edge tension, and this correlates with membrane destabilization. Indeed, charged DOPG membranes at low ionic strength were shown to display very low or even negative edge tension when pore opening occurs spontaneously ^[42]^. Under similar conditions, dimirystoyl phosphatidylglycerol (DMPG) bilayers become extensively perforated along their gel-fluid transition region ^[43]^.

The reason for the observed decreased edge tension of charged membranes is likely related to the fact that charged lipid headgroups in general have a larger effective area due to electrostatic repulsion among neighboring lipids and hydration of the charged phosphate groups ^[44]^. As a result, lipids with more conical shapes favor positive monolayer spontaneous curvature and thus help to stabilize the pore rim ^[42]^. Additionally, in cases of high surface charge density and low ionic strength, the repulsion along the pore rim was shown to render the membrane unstable upon the formation of holes ^[45]^. Regarding the stabilizing effect of calcium, theoretical predictions suggest that relatively high concentrations (≥ 5 mM) of monovalent ions such as NaCl are required for stabilization via charge screening, whereas remarkably lower concentrations of multivalent cations such as Ca^2+^ (as low as 0.1 mM) reverse the instability behavior, dramatically increasing γ, and as a result pore formation and expansion are no longer favorable ^[46]^. The effects of multivalent cations are interpreted as a result of high biding efficiency, reducing electrostatic contributions and inducing attraction between charged lipids (membrane condensation) ^[46]^. Our observations of the decrease in bursting in the presence of calcium demonstrate the combined effects of charge neutralization, membrane condensation and a suppression of curvature effects: When Ca^2+^ binds to an anionic lipid, it counteracts the lipid positive molecular curvature due to charge neutralization and in addition causes lipid condensation due to surface dehydration ^[39, 47]^. As a result, pore edge tension increases and the membrane is stabilized, which is expressed in a reduced fraction of the compromised vesicles (Figs. 2B, 3). This effect is consistent with enhanced stability of red blood cells against rupture attributed by divalent ions ^[48]^.

Membrane charges, especially when asymmetrically distributed as in cell membranes, affect membrane curvature. We hypothesize that the membrane spontaneous curvature contributes to destabilizing the porated membrane causing the extreme response of vesicle bursting. Deflation of charged electroformed vesicles containing PG has been reported to lead to tube formation ^[26]^, which is an evidence for non-zero spontaneous curvature ^[49]^. The latter was ascribed to asymmetric distribution of PG in the bilayer (resulting from the preparation approach) ^[26]^, which in terms of asymmetry renders our vesicles closer to plasma membranes. The tube suboptical diameter (< 200 nm) is consistent with that of tubes we observe after vesicle poration. The post-poration tubes also show that the membrane spontaneous curvature has not relaxed during the bursting suggesting that possible interleaflet exchange across the pore rim ^[50]^ has not suppressed the membrane asymmetry. In the following, we examine how this non-zero spontaneous curvature in addition to the lowered edge tension may contribute to vesicle destabilization.

The catastrophic effect of vesicle bursting could be understood from simple energetic considerations. The total curvature elastic energy *E* of a vesicle with a pore of radius *r* can be expressed as a sum of bending and edge energy. If the total vesicle energy decreases as the pore expands (*dE/dr* < 0), it will be unfavorable to close the pore, the vesicle will be destabilized and tend to burst. From these considerations, we speculate that edge tension contributions are dominated by curvature terms, which presumably drive the transformation of membrane area into tubes stabilized by the spontaneous curvature. It can be shown that the above energetic condition implies for the spontaneous curvature 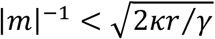 (see SI Text S3), where *κ* is the membrane bending rigidity. The edge tension of 50 mol% PG membranes is *γ* ≈ 23 pN (Fig. 4B), while the membrane bending rigidity measured in similar ionic strength conditions is *κ* ≈ 31 k_B_T ^[40c]^, where k_B_T is the thermal energy. We thus obtain that the pores will tend to expand leading to bursting for magnitude of the spontaneous curvature of the order of 1/110 nm. This value is consistent with the observation of tubes with suboptical diameter (note that for cylindrical tubes *m* ≡ − 1*/*2*R*_*cyl*_).

Although it is not clear why charged membranes containing lipids with two unsaturated tails are more prone to disruption, this may be related to the increase in membrane fluctuations due to their lower bending rigidity. High-amplitude fluctuations were shown to increase the propensity to membrane rarefactions (prepores) that may eventually evolve into a large pore due to a lower activation energy of pore formation ^[51]^.

Because the low membrane stability is rendered by charged lipids, one may wonder why bursting is not observed *in vivo*. When subject to poration (induced by electric pulses, tension, osmotic stress or membrane-active molecules), cells should exhibit similar bursting behavior as the one reported here since their membrane contains asymmetrically distributed charged lipids. Indeed, stable holes and long-term poration has been observed, see e.g. ^[10c]^. However, no full destabilization response such as whole-cell bursting is observed. Several reasons account for this: (i) The plasma membrane has rich compositional complexity and is enriched in lipids with only one saturated chain, which have a lower tendency of disruption; and lipids such as sphingomyelin and cholesterol are known to increase membrane edge tension ^[16, 23b]^. (ii) Cells are subjected to internal mechanical constraint imposed by the cytoskeleton, which stabilizes the cell membrane ^[11b, 48a]^ as shown also on model systems ^[52]^. (iii) The high ionic strength media screens possible charge-induced destabilization and the presence of extracellular calcium ions at millimolar concentration can stabilize the membrane. (iv) Cells possess a number of repair mechanisms acting on different time scales and requiring substance and/or energy supply ^[4]^. Thus, despite the presence of charged lipids, cell plasma membranes do not disintegrate even if prone to disruption presumably taking immediate advantage of their environment and composition as energetically cheaper response compared to costlier repair pathways. Instead of bursting, pores are stabilized and can either lead to cell death by lysis or resealing depending on the media. The latter is the key to the efficiency of electroporation-based protocols for drug or gene transfer in cells. For example, the stabilizing role of calcium observed here is intimately related to the success of biotechnological developments in the latter years employing calcium electroporation for cancer treatment which relies on locally increasing calcium concentration ^[6c]^. On the contrary, decreased stability of negatively charged cancer cells in electrochemotherapies might be a key to the treatment efficiency. Our results suggest that membrane charge might be an important but not yet well-characterized regulating agent in these protocols.

## 4. Conclusions

We studied the origin of bursting of charged multicomponent GUVs upon pore formation. The decreased membrane stability, which can lead to full collapse of the vesicle, is a consequence of the reduction in membrane edge tension and plausibly membrane asymmetry. During collapse, the quasi-flat membrane of the vesicle is converted into curved structures through the consumption of lipids around the pore rim, demonstrating the associated role of membrane spontaneous curvature on membrane stability. Importantly, complete vesicle disruption is only the most extreme response to destabilization among other intermediate behaviors. The results described here provide direct evidence for theoretical predictions ^[42, 46]^. The decreased stability scales with the fraction of charged lipids, is enhanced in membranes containing lipids with unsaturation in both lipid tails and the effects are partially or completely reversed by ions in the medium. Membrane asymmetry apparently plays a destabilizing role and would be interesting to explore in the future using emerging approaches for preparation of asymmetric GUVs. It would be also attractive to employ the experimental platform and approaches for pore generation and observation developed here to unravel the wound-healing mechanisms of repair proteins such as annexin and the stabilizing role of calcium ^[53]^. The fact that bursting as described here has never been observed *in vivo* demonstrates that biological membranes are largely stable against disruption and highlights how nature has created means to stabilize charged membranes, which would otherwise be unable to sustain a stable cell, and hence life.

## 5. Materials and Methods

### Materials

All materials and chemicals were used as obtained without further purification. POPC (palmitoyl-oleoyl phosphatidylcholine), DOPC (dioleoyl phosphatidylcholine), POPG (palmitoyl-oleoyl phosphatidylglycerol), DOPG (dioleoyl phosphatidylglycerol), DPPE-NBD (1,2-dipalmitoyl-*sn*-glycero-3-phosphoethanolamine-*N*-(7-nitro-2-1,3-benzoxadiazol-4-yl)), DPPE-Rh (1,2-dipalmitoyl-*sn*-glycero-3-phosphoethanolamine-*N*-(lissamine rhodamine B sulfonyl) (ammonium salt)), tail labelled NBD-PE (1-oleoyl-2-{6-[(7-nitro-2-1,3-benzoxadiazol-4-yl) amino] hexanoyl}-sn-glycero-3-phosphoethanol-amine), NBD-PG (1-oleoyl-2-{6-[(7-nitro-2-1,3-benzoxadiazol-4-yl)amino]hexanoyl}-sn-glycero-3-[phospho-rac-(1-glycerol)] (ammonium salt)) and the ganglioside GM1 were purchased from Avanti Polar Lipids (Alabaster, AL). Glucose, sucrose, NaCl, CaCl_2_, EDTA, Triton X-100 and the fluorescent dyes sulforhodamine B (SRB), calcein, CB-Dex and TR-Dex were purchased from Sigma Aldrich (St. Louis, MO USA). Atto-647 was purchased from Atto-Tec (Siegen, Germany). All lipid dyes were dissolved in chloroform and the solution was stored at stock solutions at -20°C until use. The dyes 5-Hexadecanoylaminofluorescein (5-HAD), DiIC_18_(1,1’-Dioctadecyl-3,3,3’,3’-Tetramethylindotri-carbocyanine Iodide) and the Alexa 488-labeled fragment B of cholera toxin were purchased from Thermo Fisher scientific (Waltham, MA USA). The membrane dye 5-HAD was dissolved in a mixture of chloroform:methonol (2:1 vol) and also stored at -20°C. The content dye Atto-647 dye was dissolved in a DMSO 80% volume in water and stored at 4°C.

### GUV preparation

Lipids were obtained as a powder, stock solutions were prepared in chloroform and stored under -20^°^C before use. Working solutions of 3 mM with the desired lipid mixture, containing or not a membrane dye, were prepared and used for GUVs formation. GUVs were prepared by the electroformation method ^[54]^ as described in details in ^[21]^. Briefly, ∼8 µl lipid solution was spread on a pair of conductive indium tin oxide coated glass plates and dried under a stream of nitrogen. The plates were sandwiched using a 1 mm Teflon spacer forming a chamber with ∼1.5 mL volume and connected to a function generator. An AC field of 1.7 V_pp_ (nominal voltage) and 10 Hz was applied and the chamber was filled with a 0.2 M sucrose solution at room temperature and left for 1-2 hours. For dye encapsulation inside the GUVs, the hydrating solution contained the specified dye at a concentration of 2-5 µM. When the sample contained fluorescent molecules (lipids or water-soluble dyes), growth was carried out in the dark.

### GUV observation and pulse application

Different modes of observation were employed. GUVs were observed using a Zeiss Axiovert 200 (Jena, Germany) phase contrast microscope equipped with a Zeiss AxioCam HSm camera or alternatively an Axio Observer D1 (Jena, Germany) microscope equipped with a sCMOS camera (pco.edge, PCO AG, Kelheim, Germany) for fast fluorescence recordings (up to 300 frames per second, fps) was used. For high temporal resolution acquisition (up to 10 000 fps in our experiments), a fast digital camera HG-100 K (Redlake, San Diego, CA) was used and the samples were illuminated with a mercury lamp HBO W/2. GUVs were also observed with confocal scanning fluorescence microscopy using a Leica TCS SP5 or a Leica SP8 (Wetzlar, Germany) through a 40x (0.75 NA) air or a 63x (1.2 NA) water-immersion objectives. CB-Dex was excited at 405 nm and its emission was detected at 410–480 nm. The green dyes DPPE-NBD and calcein were excited with an Argon laser line at 488 nm and emission was collected at 495-550 nm, whereas the red dyes DPPE-Rh, SRB and TR-Dex were excited with a diode-pumped solid-state laser at 561 nm and the signal was collected at 565-620 nm. The far-red dye Atto-647 was excited with a 633 laser line and its emission collected at 640-720 nm. Images were acquired in the sequential mode to avoid cross talk and snapshots were typically recorded at 512×512 pixels images and 400 Hz scanning speed in the bidirectional mode with two line averages, whereas the fast vesicle electrodeformation and poration processes were typically recorded with faster settings. Images were quantified using the Leica LAS X software (Jena, Germany) and ImageJ (NIH, USA).

For the electroporation experiments, the vesicles were diluted ∼10 fold in an isotonic glucose solution containing the desired amount of the appropriate additive and/or dye and placed in an electrofusion chamber containing two parallel cylindrical electrodes (92 mm radius) spaced by 500 µm ^[2b]^. The chamber was connected to an Eppendorf multiporator (Eppendorf, Germany), in which field strength and duration can be controlled from 50 to 300 V and 50 to 300 µs, respectively. If not otherwise indicated, vesicles were subjected to a single DC pulse (3 kV/cm, 150 □s) and the response was observed for 2-3 minutes. This procedure was repeated 3-5 times for each composition, every time on a fresh sample. To improve statistics, the quantification of vesicle responses to DC pulses was performed at lower magnification using a 10× objective (NA 0.25). We counted the number of GUVs that burst or lose contrast relative to all vesicles in the field of view. Alternatively, we measured their change in contrast over time (see e.g. Fig. S6). To record a typical event, higher magnifications (40× or 63× air objectives, NA 0.6 and 0.75, respectively) were employed.

## Supporting Information

Supporting Information is available.

## Acknowledgements

R.D. and R.B.L are thankful to R. Lipowsky, J. Steinkühler and M. Miettinen for input on vesicle stability. We thank R. Knorr and D. Duda for many fruitful discussions. R.B.L. thanks T. Robinson for the help with the edge tension analysis. The financial support of FAPESP (08/10544-0, 11/22171-6, 13/07246-5 and 16/13368-4), CAPES and INCT-FCx are acknowledged. R.B.L. and R.D. acknowledge the stimulating environment of the MaxSynBio consortium, which is jointly funded by the Federal Ministry of Education and Research in Germany and the Max Planck Society.

## Supporting Information

### Text S1. Electroporation conditions

The transmembrane potential built during the pulse is given by ^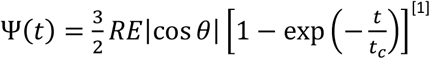 [1]^, where *R* is the vesicle radius, *E* is the field strength, *θ* is the tilt angle between the electric field and the surface normal, *t* is time and 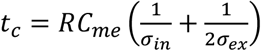 is the charging time with the membrane capacitance *C*_*me*_ and the conductivities of the solutions inside and outside the vesicle, *σ*_*in*_ and *σ*_*ex*_, respectively. Above the electroporation threshold, the transmembrane potential *Ψ* cannot be further increased and the membrane becomes permeable to ions at a certain critical transmembrane potential, *Ψ*_*c*_ ≈ 1V; see e.g. ^[2]^. In the experimental conditions explored in this work, *σ*_*in*_ ≅ 10 µS/cm, *σ*_*ex*_ ≅ 4–300 µS/cm (see also Table S1), *C*_*me*_ ≅ 0.01 F/m^2^, the transmembrane potential reaches *Ψ*_*c*_ for vesicles with radii above 7–10 µm.

**Table S1.** Conductivity values of the solutions used. Values were obtained from three individual measurements of each solution from a single batch using a conductivity meter SevenEasy (Mettler Toledo, Switzerland).

| Medium | Conductivity ($\mu S/cm$) |
| --- | --- |
| Water | $1.5 \pm 0.1$ |
| Sucrose (200 mM) | $10.1 \pm 0.2$ |
| Glucose (200 mM) | $3.9 \pm 0.1$ |
| Glucose + 1 mM NaCl | $177.0 \pm 1.7$ |
| Glucose + 1 mM EDTA | $272.3 \pm 1.1$ |
| Glucose + 0.5 mM CaCl<sub>2</sub> | $277.7 \pm 0.6$ |

**Figure S1.**
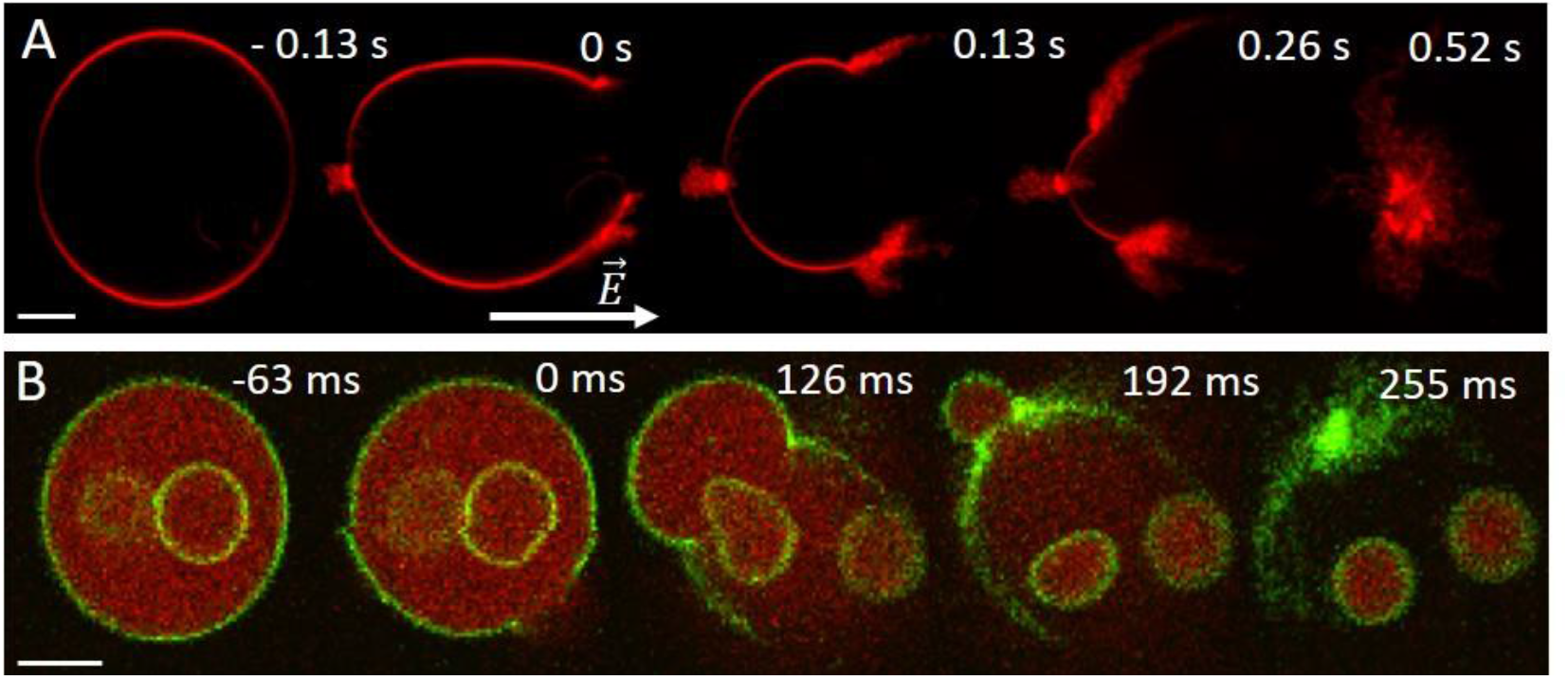
Bursting of charged GUVs (POPC:POPG; 1:1 mol) as observed in confocal microscopy cross-sections; the field direction is indicated by an arrow. (A) Upon bursting, the vesicle membrane around the pore rim is transformed into tubular structures whereby the tubulation is more pronounced at the vesicle pole facing the cathode. (B) Bursting results in fast and complete release of encapsulated content (internal vesicles and encapsulated SRB, green) from the GUVs; the same sequence is displayed in Movie S2. Scale bars: 10 µm. In (A), the membrane is labeled with 0.1 mol% DPPE-Rh, whereas in (B), the membrane contains 0.5 mol% DPPE-NBD (false color green) and the vesicle encapsulates 2.5 µM SRB (red).

**Figure S2.**
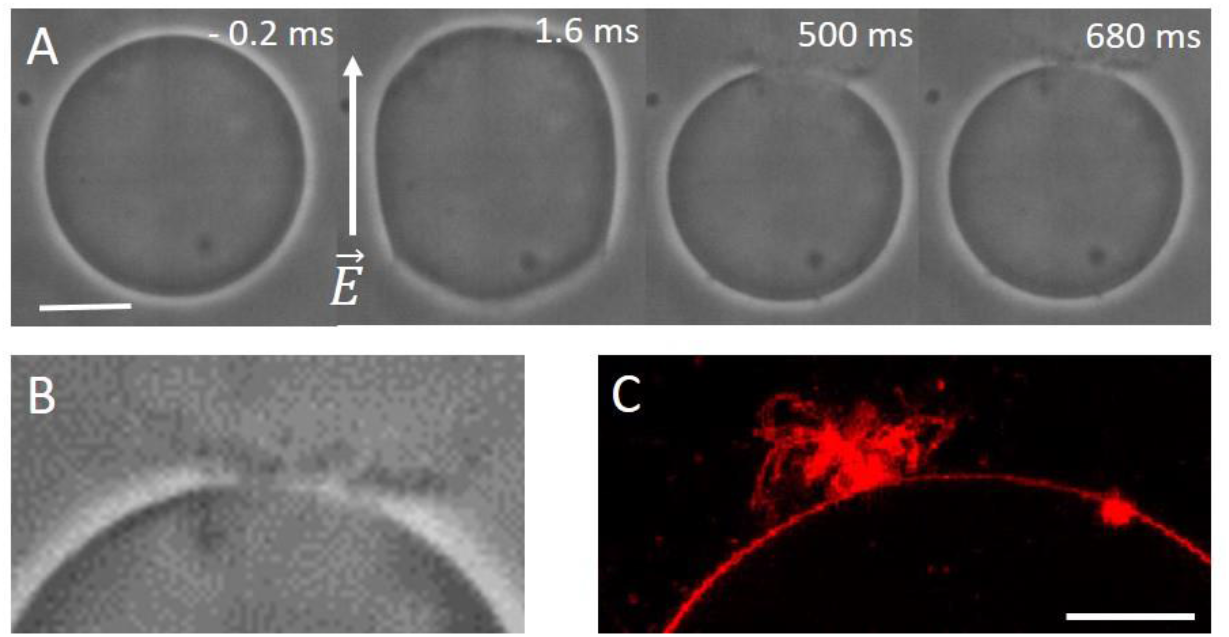
Partial bursting of charged (POPC:POPG; 1:1 mol) GUVs. (A) Phase contrast snapshots showing macropore closure associated with consumption of the membrane and conversion into tubular structures accompanied by a decrease in GUV size. The numbers correspond to time relative to the onset of the pulse. Electroporation in the presence of 1 mM NaCl. (B) Zoomed-in and enhanced image of the tubulated region in the last snapshot in panel (A). (C) Confocal cross section of another GUV showing the tubules formed in the region where the macropore closed. The GUV in (C) contains 0.1 mol% DPPE-Rh. Scale bars correspond to 20 µm in (A) and 5 µm in (B).

**Figure S3.**
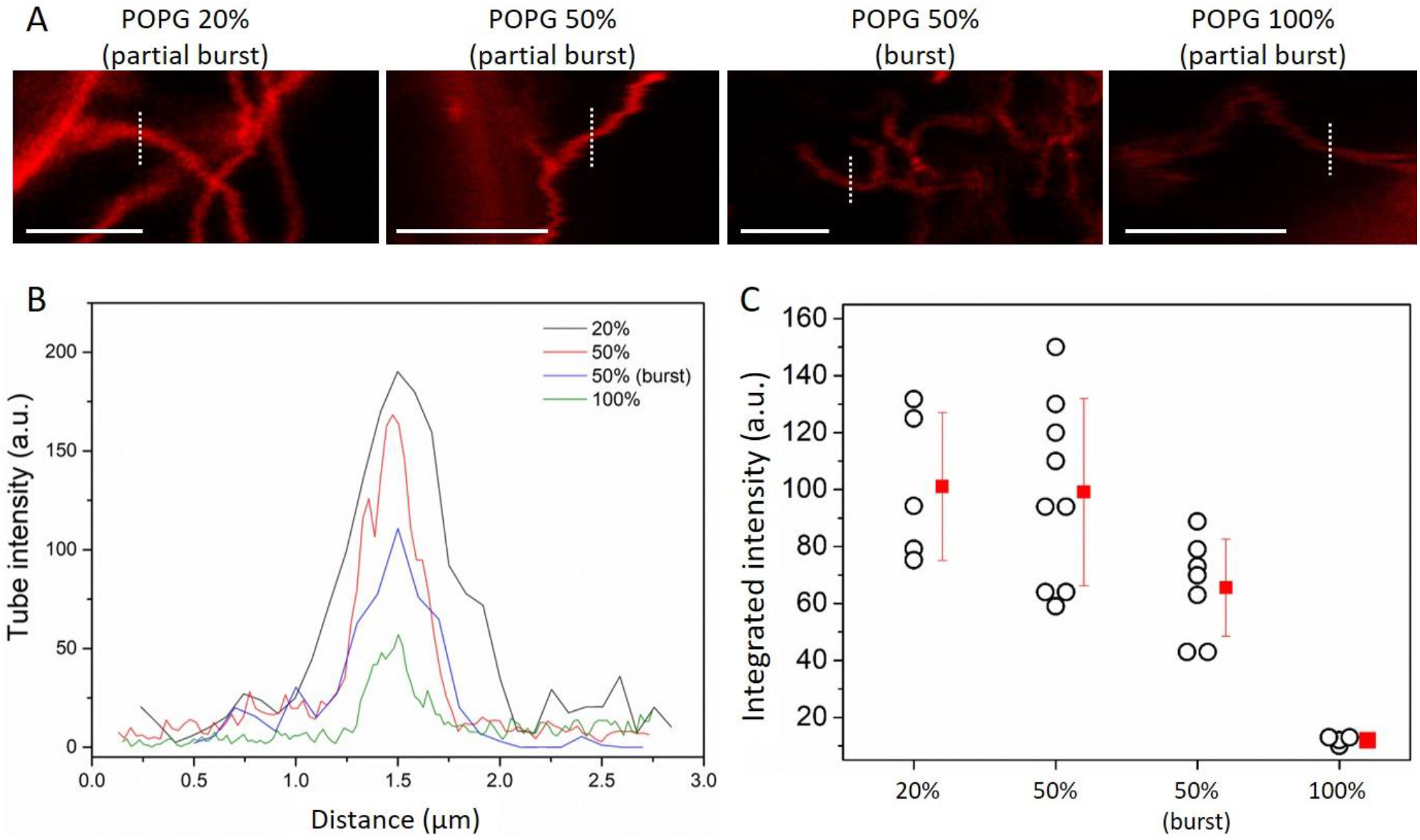
The diameter of lipid tubes formed on negative GUVs decreases with increasing membrane charge. To assess trends in the tube thickness, the tubes were imaged using identical microscope settings and a dye (DPPE-Rh) that displays no curvature preference ^[3]^. (A) Images of lipid tubes formed after complete vesicle bursting (burst) or partial vesicle bursting (partial burst) where the tubes remain attached to the mother GUV. The POPG fraction in the membrane varies from 20 to 50 to 100 mol% from left to right. The images show weaker intensity associated with thinner tubes as the PG fraction increases. Scale bars: 5 µm. (B) Tube fluorescence intensity profiles from the vertical lines shown in (A). Line profiles were always taken in the same direction to minimize polarization issues. (C) Area-integrated peak intensities measured on tubes formed upon bursting (burst) or still attached to partially burst GUVs for increasing POPG mol%. Each open circle corresponds to a measurement on an individual tube. Mean and standard deviation are also shown (red). The decrease in intensity is associated with decreased tube diameter.

**Figure S4.**
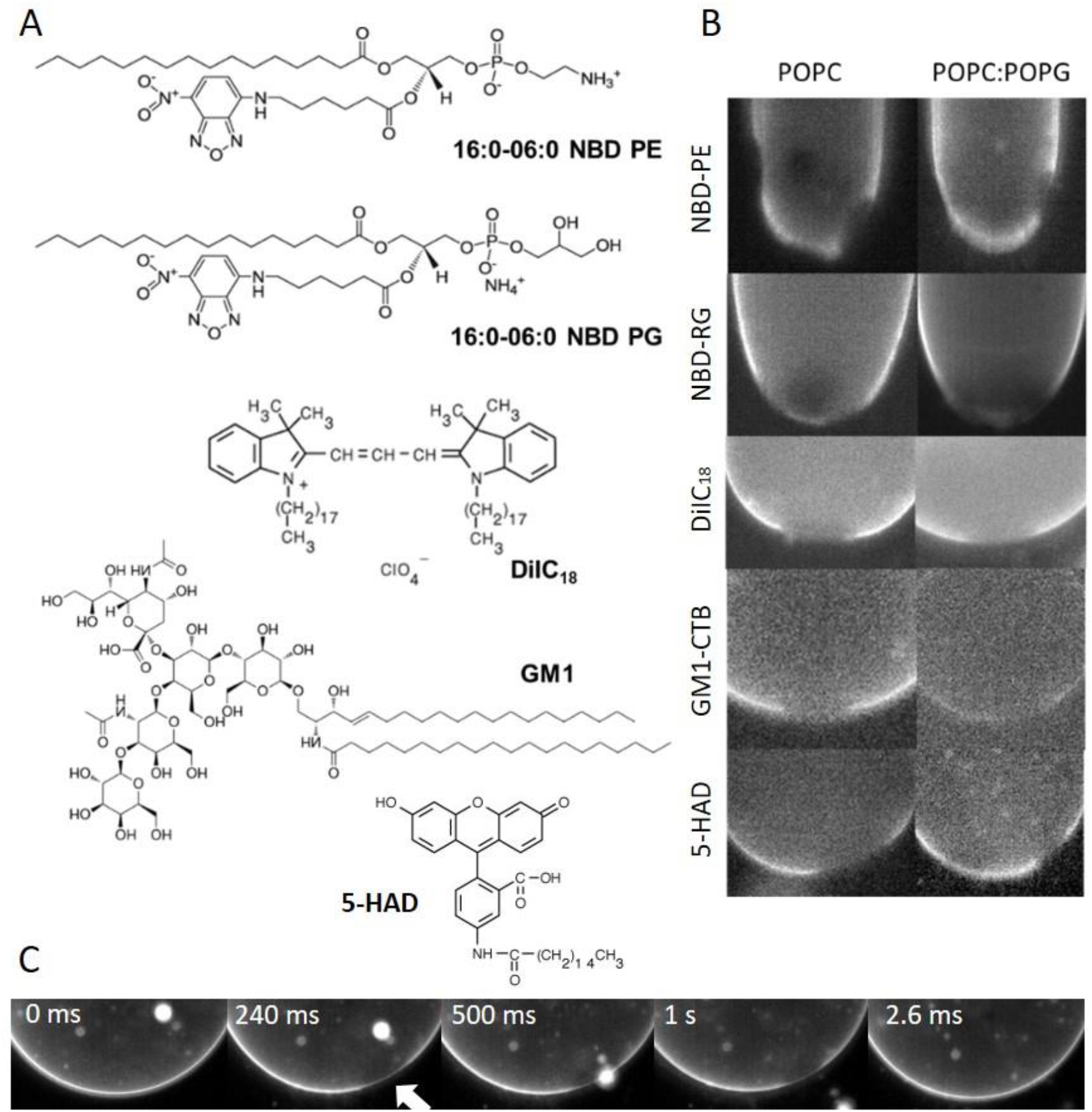
Composition in the pore rim vicinity is similar to that of regions of intact membranes as viewed with fluorescence markers. (A) Chemical structure of (some of) the used fluorescent probes of different charge and molecular geometry. Their concentrations in the membrane was varied from 0.1 to 1 mol%. (B) Snapshots of pores in neutral (POPC) or negative (POPC:POPG) GUVs doped with different dyes: neither accumulation nor depletion of the probes of diverse biophysical properties can be detected. GUV sizes varied from 30 to 60 µm. The individual frame exposure time was 5 ms or 20 ms. (C) Snapshots of the closure of long-lasting macropore on a negative GUV. Note the exit of internal structures through the pore. The membrane is labeled with NBD-PG.

**Figure S5.**
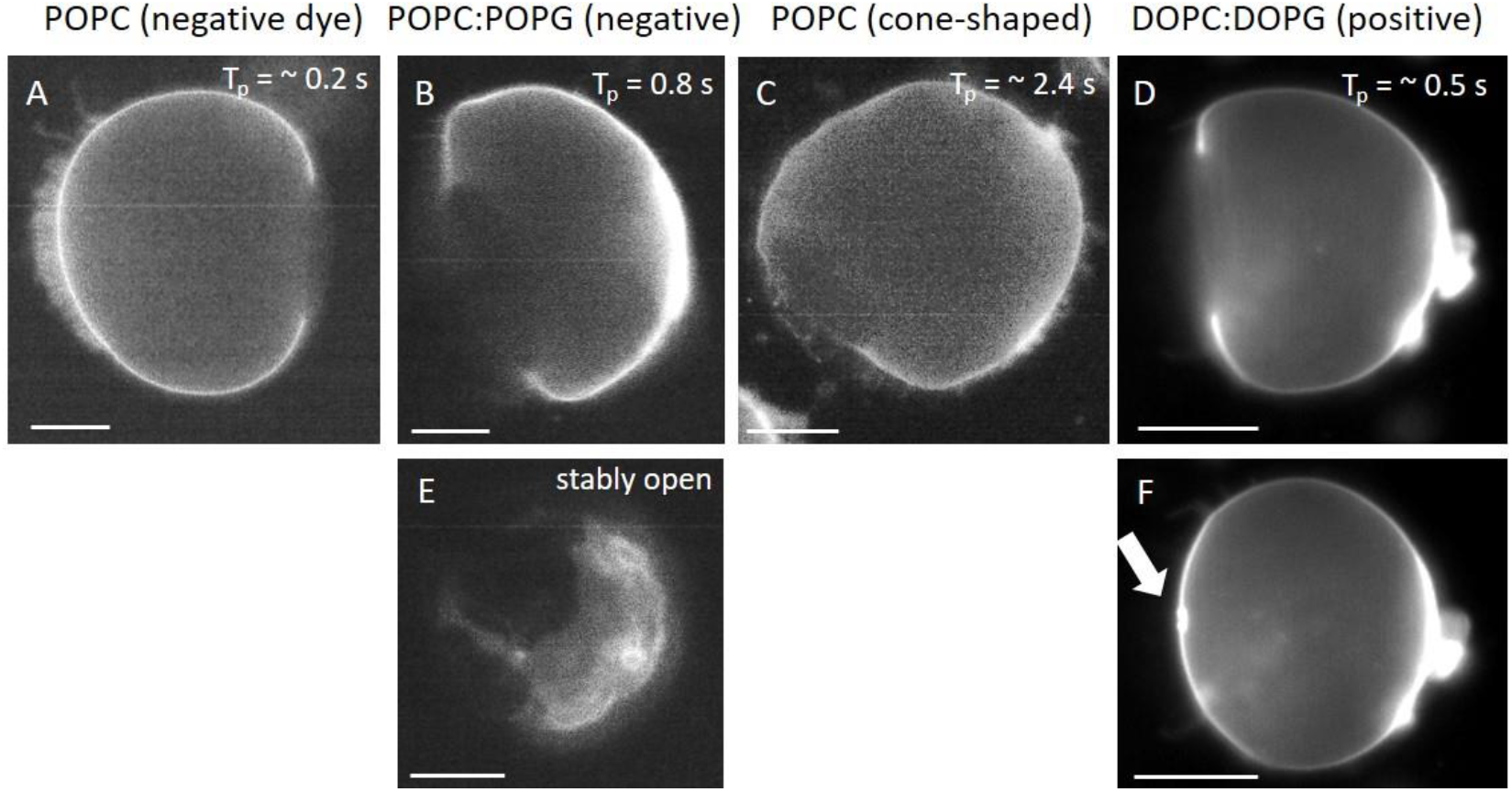
Lipid analogues do not accumulate in the rim of long-lasting pores. Following the protocol in ^[4]^, GUVs were immobilized in 1% (w/v) isotonic agarose gel to slow down pore closure and were exposed to a single DC pulse (4 kV/cm, 8 ms). Membrane compositions and dye properties are indicated above the images. The text in brackets describes the charge and shape of the fluorescent lipid analogues used. Negative: NBD-PG; cone-shaped: 5-HAD; positive: DPPE-Rh, see corresponding structures in Fig. S4A. (A-D) Representative behavior of electroporated GUVs of different composition. The numbers (top right) show the pore lifetime (Tp) for these specific vesicles. (E) POPC:POPG 1:1 GUV with stably open pores (Tp > 1 minute). (F) Small buds formed in the region where the macropore closed (arrow). The non-spherical GUV shapes result from the confining agarose gel mesh around the vesicles. Scale bars: 20 µm.

**Figure S6.**
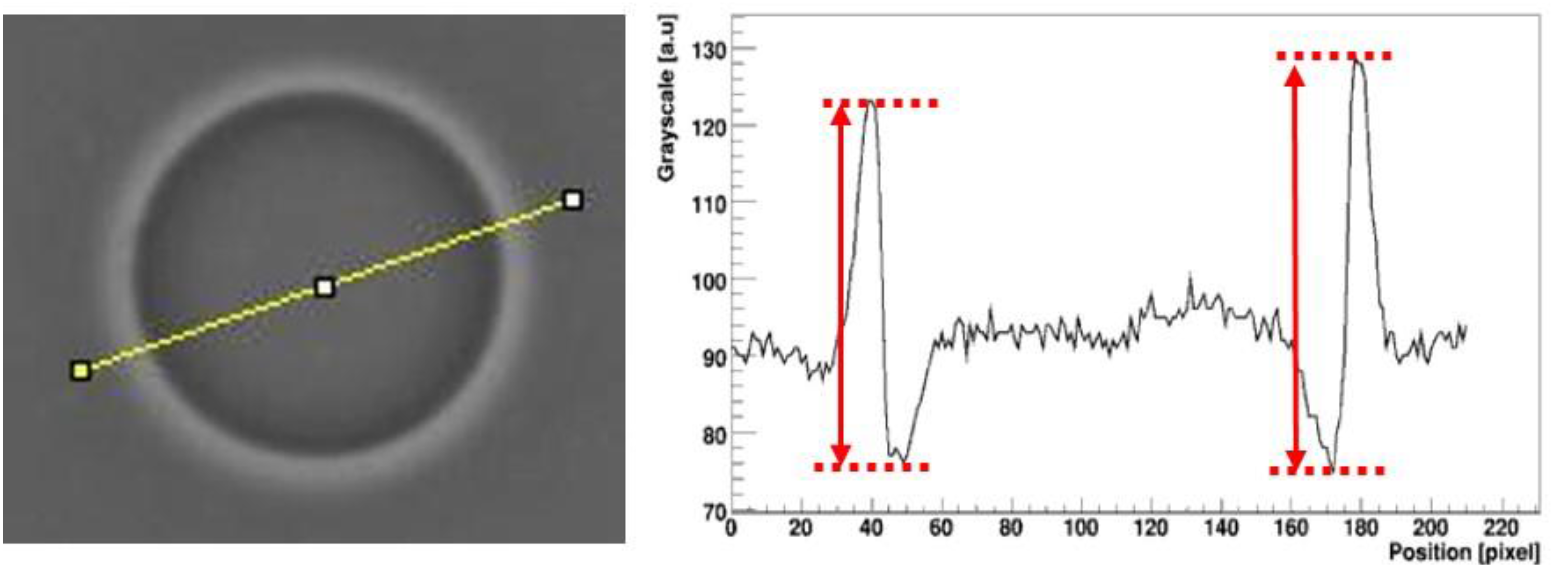
Measuring the optical contrast of a GUV imaged in phase contrast mode. A line is drawn across the vesicle (left) and the grayscale profile along this line is obtained (right). The optical contrast is defined as the average of the two heights indicated in red in the figure. The analyses were done with ImageJ.

**Figure S7.**
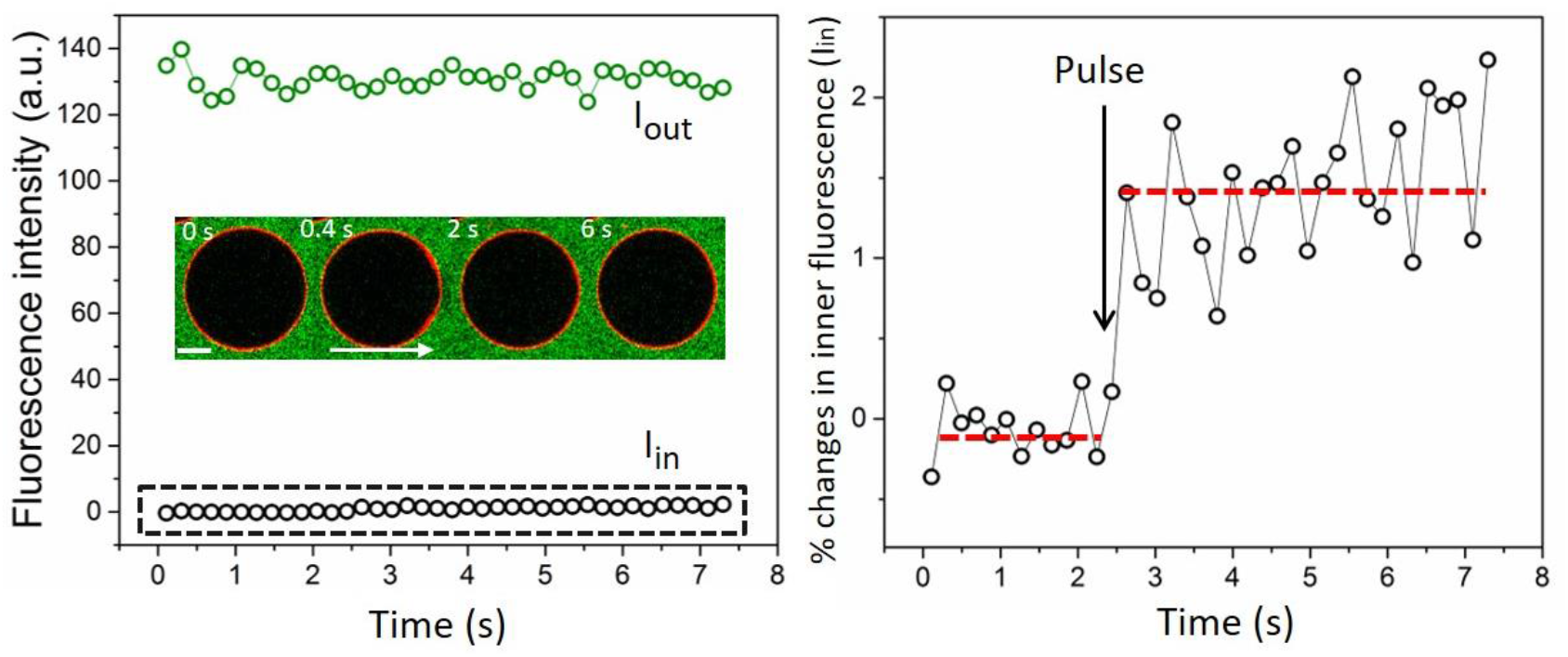
Only a small fraction of dyes enters the vesicles directly through macropores demonstrated from following the entry of a small external dye (calcein) upon GUV macroporation after which the vesicle reseals. (A) Electroporation of a negative GUV (0.1 mol% DPPE-Rh, red) in the presence of 5 µM calcein (green). The field direction is indicated with an arrow. The numbers correspond to the time after applying the pulse (3 kV/cm, 300 µs). Scale bar: 20 µm. Calcein fluorescence outside, Iout, and inside the GUV, Iin, is shown in the graph. (B) The increase in calcein fluorescence inside the GUV upon pulse application (arrow) is in the order of only ∼ 1% (same data as in A).

**Figure S8.**
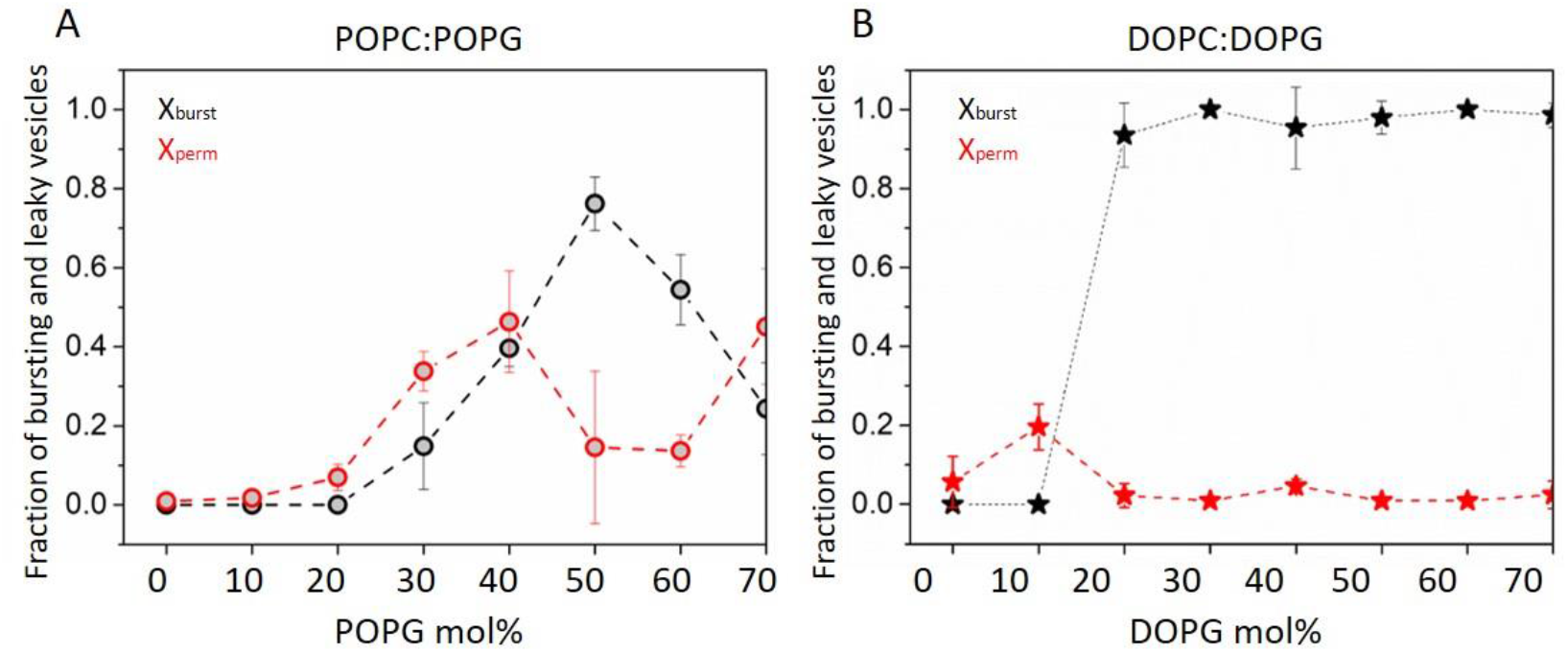
Fraction of bursting vesicles (black data), X_burst_ = n_burst_/n_GUVs_, and fraction of surviving vesicles which lose contrast (red data), X_perm_ = n_perm_/(n_GUVs_ - n_burst_), for increasing molar fractions of charged lipids. Panels (A) and (B) respectively show data for POPC/POPG and DOPC/DOPG vesicles with increasing fractions of the PG lipid. Average and standard deviation from a number of measurements on different vesicles (> 15) of a given composition are shown.

### Text S2: Edge tension measurements

Pore dynamics in vesicles follows four well-defined stages; (i) quick opening, (ii) maximum size stage that is relatively stable, (iii) slow closure, limited by leak-out, and (iv) rapid closure as theoretically modeled in ^[5]^. According to this model, the slow closure (third stage) of a circular pore of radius *r* in a GUV of radius *R*, can be directly related to the edge tension through *R*^2^*ln*(*r*) = −(2*y/*3*πχ*)*t* + *C*, where *η* is the medium viscosity (*η* = 1.133×10^−3^ Pa.s for the conditions studied here), *t* is time and *C* is a constant. *R* is assumed roughly constant during pore closure and measured after resealing of the macropore(s). The edge tension *γ* is directly calculated from the slope of the linear dependence of *R*^2^*ln*(*r*) with time, see also Fig. S9. Vesicles with radius deviation larger than 5 % were discarded for GUVs with resealing pores. The obtained mean edge tension values and standard deviations are given in Table S2. In the case of partial bursting where the vesicle size decreases as the pore closes (data in magenta in Fig. 4), the edge tension was evaluated considering the initial and final vesicle radius and averaged (error bars in Fig. 4 show the corresponding deviations resulting from vesicle size changes).

**Table S2.** Edge tension values obtained for different membrane compositions.

| Composition | Additives | $\gamma$ (pN) |
| --- | --- | --- |
| POPC | 0.5 mM NaCl and 0.2 mM EDTA | $39.5 \pm 5.2$ |
| POPC with 8 mol% POPG | 0.5 mM NaCl and 0.2 mM EDTA | $40.8 \pm 5.1$ |
| POPC with 16 mol% POPG | 0.5 mM NaCl and 0.2 mM EDTA | $39.5 \pm 4.5$ |
| POPC with 24 mol% POPG | 0.5 mM NaCl and 0.2 mM EDTA | $40.8 \pm 7.4$ |
| POPC with 30 mol% POPG | 0.1 mM NaCl | $40.8 \pm 10.2$ |
| POPC with 35 mol% POPG | 0.1 mM NaCl | $39.9 \pm 8.6$ |
| POPC with 42 mol% POPG | 0.5 mM NaCl and 0.2 mM EDTA | $31.3 \pm 6.1$ |
| POPC with 50 mol% POPG | 0.5 mM NaCl and 0.2 mM EDTA | $23.4 \pm 6.0$ |
| POPC with 50 mol% POPG | 1 mM $\text{CaCl}_2$ | $62.1 \pm 11.5$ |

**Figure S9.**
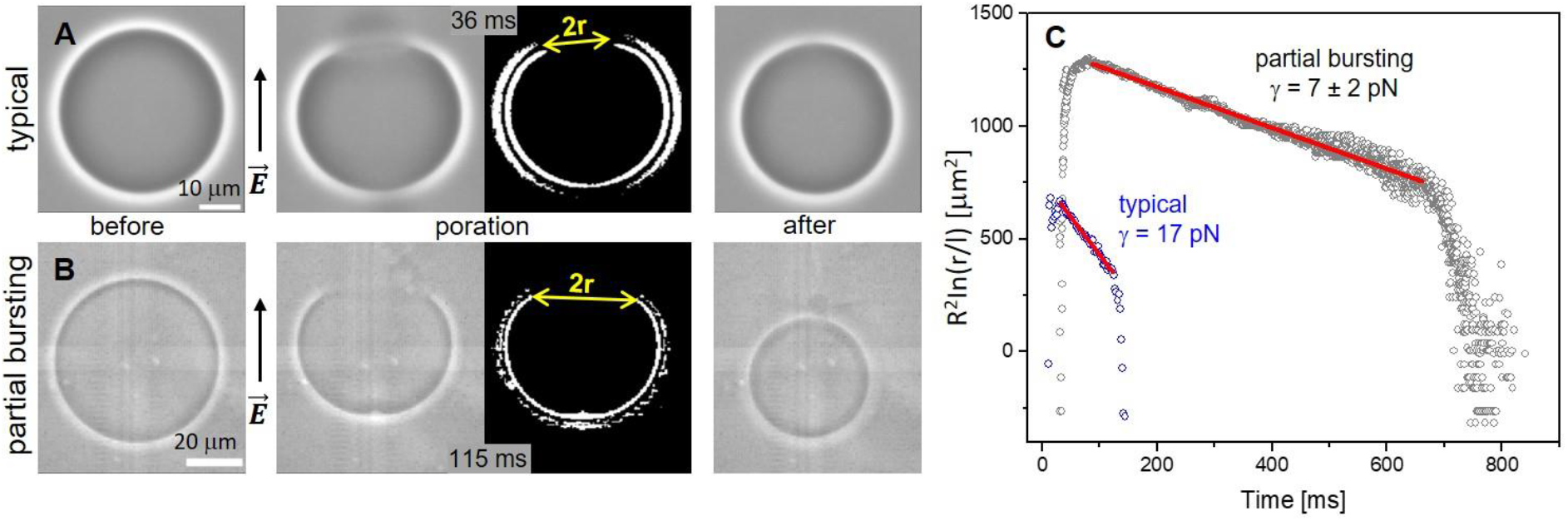
Pore edge tension measurement for negative POPC:POPG (1:1) GUVs. (A, B) Phase contrast snapshots of a GUV before, during and after pore closure, respectively, for typical pore closure (upper row) and partial bursting cases (lower row). The third snapshots in the rows are processed images (same as the second snapshots) used for measuring the pore radius *r*. The processing of the phase contrast images involves background subtraction and binarization as introduced in ^[6]^. (C) Edge tension measurements showing the pore opening and closure and the fit for the slow closure stage (red lines) according to *R*^2^*ln*(*r*) = −(2*y/*3*πχ*)*t* + *C* for typical (blue data) and partial bursting cases (black data). In the graph, the pore size *r* in µm is rescaled by *l* = 1 µm. Pore closure is significantly slowed down for the case of partial bursting. Accordingly, the edge tension extracted from the vesicle with a typical macropore is 17 pN, while the one obtained from the vesicle exhibiting partial bursting is only 7 ± 2 pN.

**Figure S10.**
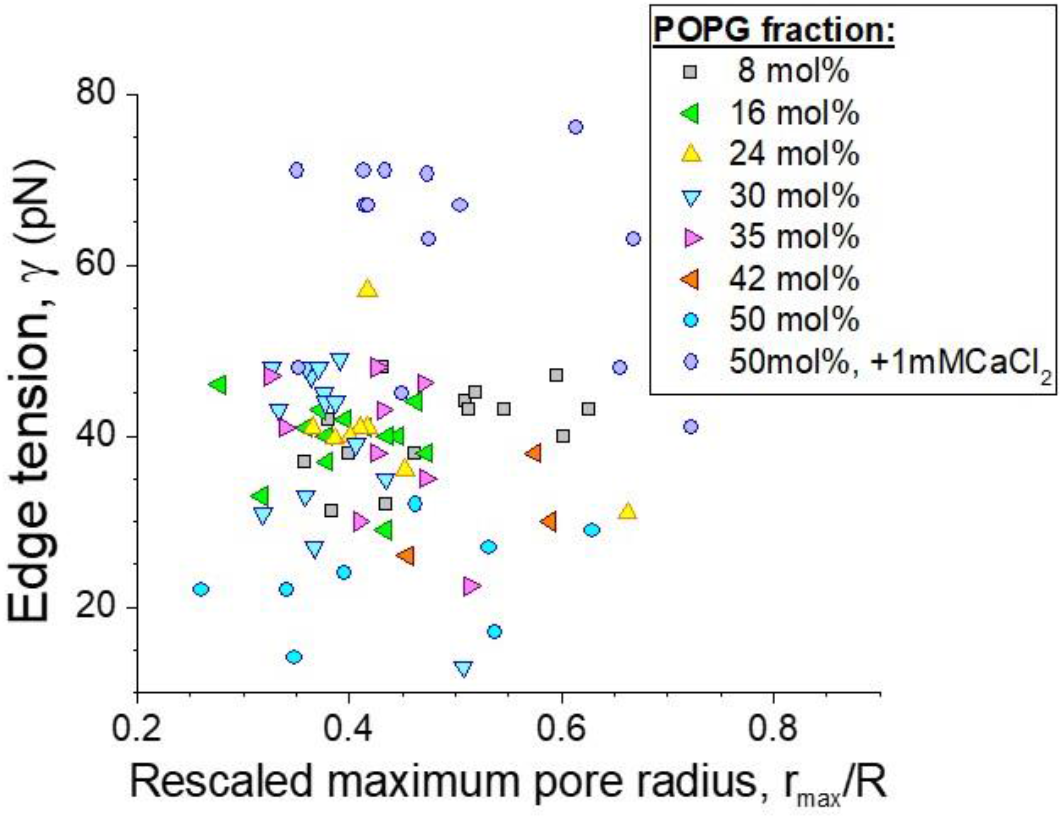
The measured edge tension does not depend on the maximum pore size rmax (here, rescaled by the GUV size, R) as measured for all POPC:POPG compositions in the absence and in the presence of 1 mM CaCl2.

### Text S3. Energetic considerations for pore expansion and closure

**Figure S11.**
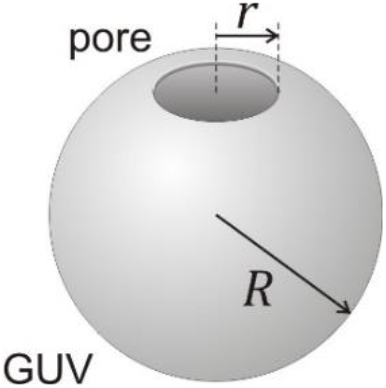
Sketch of a vesicle with a pore. The vesicle radius, *R*, and pore radius, *r* are indicated.

The energy of a porated vesicle of radius *R* is a sum of the Helfrich and the rim energy:

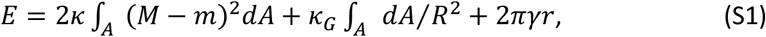

where *κ* is the membrane bending rigidity, *M* = 1*/R* is the vesicle mean curvature, *m* is the membrane spontaneous curvature, *κ*_*G*_ is the Gaussian curvature modulus, *r* is the pore radius and *γ* is the edge tension. The integral in Eq. S1 is over the vesicle (nonporated) area 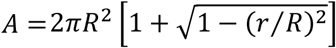.

For a pore to have the tendency to expand, as is the case in bursting PG-containing vesicles, the energy of the system should decrease, i.e.

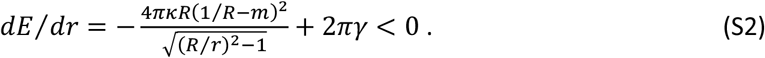

Above we assumed that the vesicle radius remains constant, which for small changes in the pore radius is indeed the case. We see that the Gaussian curvature contribution (second term in Eq. S1) cancels out, which is understandable because the topology of a vesicle with expanding pore is preserved. Equation S2 implies that the edge tension term is dominated by curvature contributions.

PG-containing electroformed vesicles exhibit inward tubes ^[7]^ with diameter below the resolution of confocal imaging, 2*R*_*cyl*_ < 200nm. The associated spontaneous curvature for inward cylindrical tubes is *m* ≡ − 1*/*2*R*_*cyl*_ implying that 1*/R* ≪ *m* and can be ignored. The inequality S2 can then be transformed into

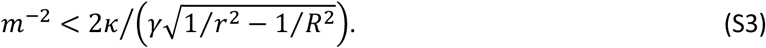

Considering that the pore radius reached at the end of the pulse is of the order of 1 µm, i.e., *r* ≪ *R*, the above expression roughly reduces to

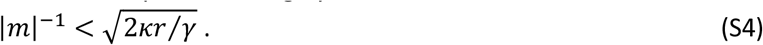

For 50 mol% PG membranes, we have *γ* = 23 pN (see data in Fig. 4B) and *κ* = 31 kBT ^[8]^. We thus obtain that the diameter of tubes stabilized by spontaneous curvature at which the vesicles tend to burst should be smaller than 105 nm. This result is consistent with the subopical tube diameters we observe upon poration (Fig. S3). The predicted relatively high spontaneous curvature is plausibly generated by the asymmetric distribution of PG in the membrane resulting from the vesicles preparation method [7].

As a consistency check, for pure PC membranes where the pores tend to close, the energy of the systems should satisfy the opposite condition *dE/dr* < 0, i.e. it is energetically unfavorable for the pores to open. These single-component PC membranes are intrinsically symmetric and exhibit zero spontaneous curvature *m* ≈ 0, which from Eq. S2 implies that 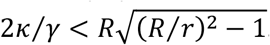. Taking for *γ* = 40 pN (Fig. 4B) and *κ* ≈2.1×10^−19^ J ^[9]^, we see that this condition is indeed satisfied.

## Supplementary movies

**Movie S1**. Bursting of a charged GUV upon electroporation observed under phase contrast microscopy. The GUV composed of POPC:POPG (1:1) in sucrose/glucose (in/out) solution was electroporated using a pulse of 3 kV/cm magnitude and 150 µs duration. The formed macropore expands, leading to a full collapse of the vesicle, leaving only tubular membrane fragments after bursting. The time stamps correspond to the initial observation time. The GUV was electroporated at time 34 ms. Vesicle diameter is around 25 µm.

**Movie S2**. Bursting of a charged GUV upon electroporation observed under epifluorescence microscopy. The GUV composed of POPC:POPG (1:1) prepared in sucrose/glucose (in/out) solution and labelled with 0.1 mol% DPPE-Rh was electroporated (3 kV/cm, 150 µs). The formed macropore expands by converting the nearly flat membranes into tubular lipids structures. The time stamps correspond to the initial observation time. The GUV was electroporated at time 17 ms. Vesicle diameter is around 30 µm.

**Movie S3**. Bursting of a charged GUV upon electroporation observed under confocal microscopy. The GUV composed of POPC:POPG (1:1), labelled with 0.5 mol% DPPE-NBD (green), prepared in sucrose/glucose (in/out) solution and encapsulating 2.5 µM SRB (red) was electroporated (3 kV/cm, 150 µs). The GUV contains a couple of smaller vesicles inside. Bursting leads to the rapid release of the encapsulated content. The time stamps correspond to the initial observation time. The GUV was electroporated at time 190 ms. Vesicle diameter is around 30 µm.

**Movie S4**. Long-lasting macropore in a charged GUV observed under epifluorescence microscopy. The GUV composed of POPC:POPG (1:1) containing 0.1% DPPE-Rh was electroporated (3 kV/cm, 150 µs) and the formed macropore remained opened for longer than 2 s. Such a long pore lifetime allowed the release of encapsulated smaller vesicles inside the GUV. Note that the fluorescence around the pore rim is homogenous, suggesting neither enrichment nor depletion of lipids. The time stamps correspond to the initial observation time. The GUV was electroporated at time 60 ms. Vesicle diameter is around 50-60 µm.

**Movie S5**. Electroporation of a neutral GUV (POPC) labelled with 0.5 mol% NBD-PG and immobilized in 1 wt% agarose. A single DC pulse (4 kV/cm, 8 ms) was applied at time ∼1.1 s. The GUV was observed under epifluorescence microscopy. Vesicle size is around 40 µm.

**Movie S6**. Bursting of a charged GUV, POPC:POPG (1:1), induced by detergent and observed at high temporal resolution under phase contrast microscopy. A concentrated solution of Triton X-100 was added to the chamber (to a final concentration of 1 mM) containing POPC:POPG (1:1) GUVs. The movie is slowed down. The time stamps correspond to the initial observation time. The macropore appears at time 12.5 ms. The expansion of the bursting membrane is very fast and is completed in less than 3 s. Vesicle size is around 50-60 µm.

**Movie S7**. Bursting of a charged GUV, POPC:POPG (1:1), induced by detergent observed under epifluorescence microscopy. A concentrated solution of Triton X-100 was added to the chamber (to a final concentration of 1 mM) containing POPC:POPG (1:1) GUVs labelled with 0.1 mol% DPPE-Rh. At initial times, Triton X-100 leads to an increase in GUV are (the membranes look floppy). The numbers correspond to the initial observation time. The macropore appears at 3.7 s and leads to vesicle bursting with dynamics identical to that induced by electroporation. At later times, the membrane fragments formed after bursting are solubilized. Vesicle size is around 50-60 µm.

**Movie S8**. Formation of submicron-sized pores before vesicle bursting. A POPC:POPG (1:1) GUV labelled with 0.5 mol% DPPE-NBD and in the presence of 2.5 µM of SRB was incubated with Triton X-100 (1 mM final concentration) and observed under confocal microscopy. The movie is sped up. The vesicle becomes permeable to SRB (at ∼ 40 s) before it bursts (at ∼3 min). The macropore formation is too fast to be observed with confocal microscopy. Vesicle size is around 30 µm.

**Movie S9**. Macropores formed on charged GUVs are stabilized in the presence of CaCl2. A POPC:POPG (1:1) GUV labelled with 0.1 mol% DPPE-Rh and in the presence of 3.5 mM CaCl2 was incubated with Triton X-100 (1 mM final concentration) and observed under epifluorescence microscopy. The detergent-induced macropore remains open for many seconds until complete GUV solubilization, allowing the exit of encapsulated vesicles. The GUV becomes dimmer over time resulting from the combined effect of membrane solubilization and DPPE-Rh photobleaching. The initial vesicle size is around 50-60 µm.

